# Partial endothelial-to-mesenchymal transition (EndMT) contributes to lumen re-organization after carotid artery ligation

**DOI:** 10.1101/2021.08.13.456319

**Authors:** Yoshito Yamashiro, Karina Ramirez, Kazuaki Nagayama, Shuhei Tomita, Yoshiaki Kubota, Hiromi Yanagisawa

## Abstract

Endothelial-to-mesenchymal transition (EndMT) is a fundamental process in vascular remodeling. Carotid artery ligation is commonly used for induction of neointima formation and vessel stenosis; however, the precise regulatory mechanism of vascular remodeling is not entirely understood. In this study, we showed that resident endothelial cells (ECs) are the origin of neointima cells and ECs transiently expressed CD45 in the early stage of neointima formation accompanied by increased expression of EndMT markers. In vitro, CD45-positive EndMT was induced by stabilization of HIF-1α with cobalt chloride or VHL inhibitor in human primary ECs, which mimicked the hypoxic condition of ligated artery, and promoted the formation of integrin α11-SHARPIN complex. Notably, a CD45 phosphatase inhibitor disrupted this complex, thereby destabilizing cell-cell junctions. These results suggest that the CD45 activity is required for the retention of an EC phenotype and cell-cell junctions during EndMT (termed “partial EndMT”). We thus propose a novel mechanism of partial EndMT that contributes to lumen re-organization during vascular injury.

## Introduction

Vascular remodeling is essential for the maintenance of vessel wall homeostasis and involves the functional and structural changes in the cells and extracellular matrix (ECM) in response to various stimuli^1^. During vascular remodeling, vascular endothelial cells (ECs) and mesenchymal cells, such as smooth muscle cells (SMCs) and fibroblasts, undergo adaptation to environmental changes and participate in the repair of the vessel wall. Integrins are a family of heterodimeric receptors that cross the plasma membrane and act as biomechanical sensors of the microenvironment^2^, linking the ECM and the cell cytoskeleton. Activation of the integrins during vascular remodeling induces intracellular signaling pathway(s), thereby changing cellular behaviors and functions.

We have previously reported that the matricellular protein thrombospondin-1 (Thbs1) is a critical component of vascular remodeling^3, 4^, especially in mechanotransduction, and controls the activation of Yes-associated protein (YAP) by binding to integrin αvβ1^5^. In vascular stenosis (neointima formation), activation of the Thbs1/YAP signaling pathway contributed to the proliferation of neointima cells, and deletion of Thbs1 in mice attenuated neointima formation upon carotid artery ligation^5^. Recently, we and others have reported that some neointima are formed without the contribution of medial SMCs, suggesting a diverse origin of the neointima cells^6, 7^. However, little is known about the initiation of neointima formation and the key regulatory processes involved in response to the cessation of blood flow.

Endothelial to mesenchymal transition (EndMT) is characterized by the loss of endothelial markers, such as vascular endothelial (VE)-cadherin and CD31 (also known as PECAM), and the acquisition of mesenchymal properties; loss of cell-cell junctions, apical-basal polarity, and the ability to form tubes; and the induction of mesenchymal markers, including alpha-smooth muscle actin (αSMA), fibroblast-specific protein-1 (FSP-1), collagen type I, and fibronectin^8, 9^. EndMT plays critical role in embryonic development and cardiovascular diseases^10-12^ such as pulmonary arterial hypertension (PAH)^13^, cardiac fibrosis^14^, and atherosclerosis^15, 16^. In addition, the process of angiogenic sprouting has been linked to EndMT^17^, but it differs from the normal EndMT process; endothelial marker(s) expression and cell-cell junctions are retained during migration and sprouting. Moreover, protein tyrosine phosphatase receptor type C (CD45)-positive endothelium has been observed in mitral valves at 6 months post-myocardial infarction^18^, and this process also preserves the endothelial phenotype undergoing EndMT. Therefore, these EndMT processes have been termed “partial EndMT”^19, 20^. Interestingly, this concept is consistent in other cell types undergoing epithelial-to-mesenchymal transition (EMT), such as mammary and pancreatic epithelial cells. Shamir et al. reported that Twist1-induced EMT, which is a critical step in metastasis, required E-cadherin expression^21^, and Liu et al. showed that overexpression of E-cadherin did not prevent in vitro invasion of cells undergoing EMT^22^. A recent study by Na et al. supported these observations and revealed that the functional activity of E-cadherin controls metastatic spreading in cells undergoing EMT^23^. Collectively, these findings suggest that partial EMT/EndMT is functionally different from normal EMT/EndMT. However, CD45 is not normally expressed in ECs, and the details of its regulatory mechanism, whether ECs undergo a normal or partial EndMT, and the exact functional relationship between partial EndMT and CD45 are not fully understood.

Here we report a contribution of partial EndMT to lumen re-organization after carotid artery ligation. We demonstrate that the neointima cells are derived from residential ECs and CD45 phosphatase (PTPase) activity is required to maintain cell-cell junctions undergoing EndMT.

## Results

### SMC- and EC-specific markers are detected in the developing neointima

Several studies have reported that neointima formation after the carotid artery ligation is associated with the de-differentiation of mature SMCs, the differentiation of circulating hematopoietic progenitor cells or activated leukocytes^24-27^. Strong expression of αSMA, a marker of immature SMCs, was consistently observed in the developing neointima at 2 and 3 weeks after carotid artery ligation (Fig. 1a, b). At 2 weeks after ligation, the EC-specific marker CD31 was also expressed in neointima cells at proximal lesions, but the mature SMC marker smooth muscle myosin heavy chain (SM-MYH) was not expressed in these cells (Fig. 1c). More evidently, as shown in Fig. 1d with high magnification, most neointima cells showed PECAM expression at 2 weeks after ligation. At 3 weeks after ligation, CD31 expression was still prominent in the neointima cells and expanded to distal lesions (Fig. 1e), although its expression level was decreased compared with that observed at 2 weeks (Fig. 1f). At 4 weeks after ligation when the neointima was established, αSMA was consistently expressed, while PECAM expression was confined to the spot-like regions and innermost cell lining (Fig. 1g). Furthermore, we examined the leukocyte markers CD11b (also known as integrin alpha M) for monocytes, CD68 for active macrophages and PDGFRα for adventitial fibroblasts. CD11b and CD68 were detected at lower levels in the developed neointima (Fig. 1h). Although PDGFRα-positive cells were detected in the developed neointima, they were not the main resource of neointima cells (Extended data Fig. 1). These results show that the neointima cells expressed both mesenchymal and EC-specific markers and suggest the possible contribution of ECs to neointima formation upon carotid artery ligation.

**Figure 1.**
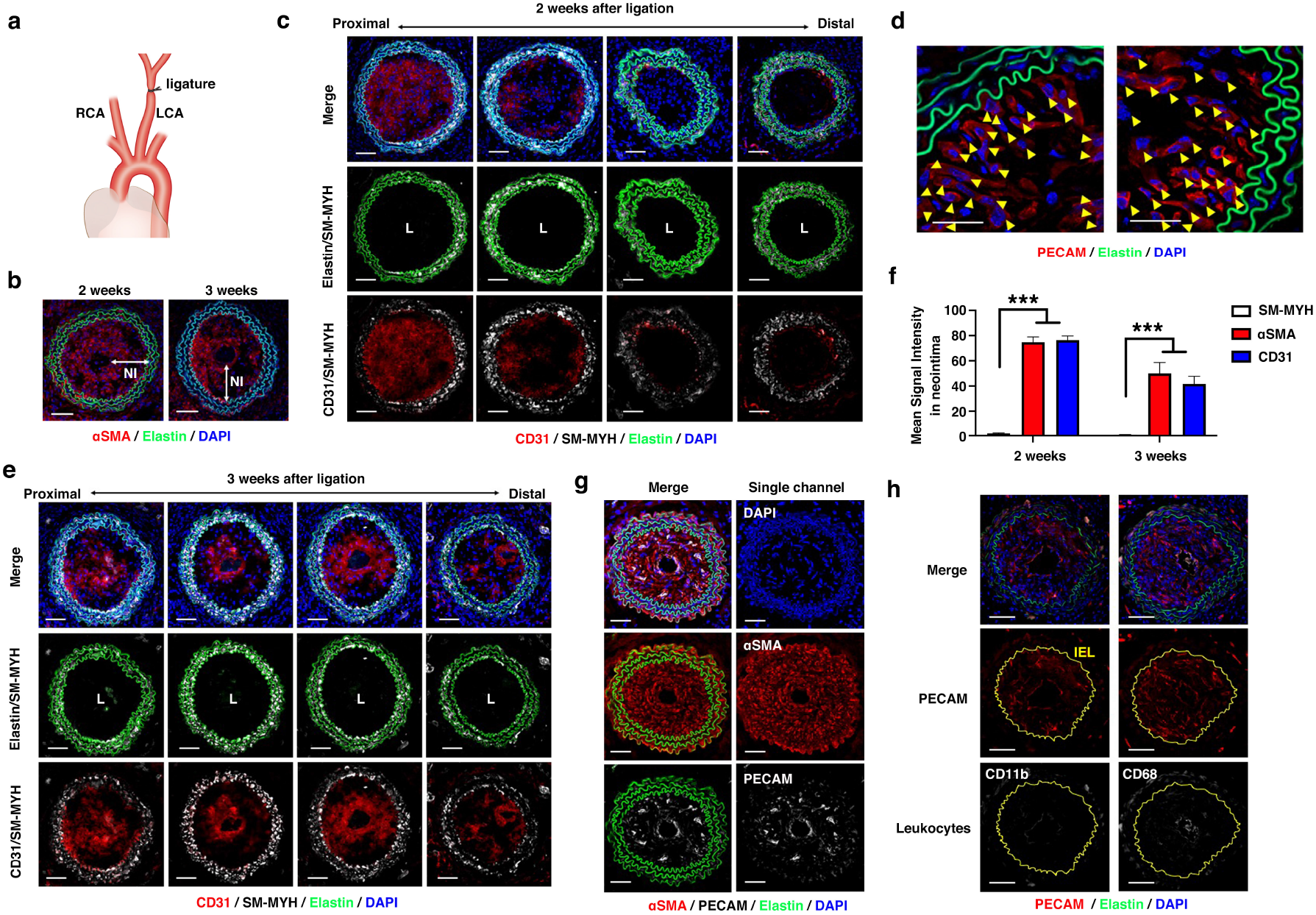
EC-specific marker CD31 and SMC-specific marker αSMA are expressed during neointima formation upon carotid artery ligation. **(a)** Diagram of left carotid artery (LCA) ligation and contralateral right carotid artery (RCA) without ligation. Image was adapted from reference^5^ with permission from the National Academy of Sciences. **(b)** Immunostaining of cross sections of ligated arteries with αSMA (red) at 2 weeks (n=5) and 3 weeks (n=4) after ligation in WT mice. NI: neointima. **(c and e)** Immunostaining of serial cross sections of ligated arteries with CD31 (red) and SM-MYH (white) at 2 weeks (in c, n=5) and 3 weeks (in e, n=4) after ligation in WT mice. Images are shown in proximal to distal direction from the ligature point. **(d)** Immunostaining of cross sections of ligated arteries with PECAM (red; yellow arrowheads) at 2 weeks (n=5) after ligation in WT mice. **(f)** Quantification of mean signal intensity of SM-MYH, αSMA, and CD31 in c and e. Bars indicate mean ± SEM. \*\*\**P*< 0.001, two-way ANOVA. **(g)** Immunostaining of cross sections of ligated arteries with αSMA (red) and PECAM (white) at 4 weeks (n=4) after ligation in WT mice. **(h)** Immunostaining of cross sections of ligated arteries with PECAM (red) and CD11b or CD68 (white) at 4 weeks after ligation in WT mice (n=4). Yellow line shows internal elastic lamina (IEL). **(b-e, g, and h)** All scale bars equal 50 μm. DAPI (blue) and autofluorescence of elastin (green) are also shown.

### Endothelial lineage tracing during neointima formation

Technological advances in genetically engineered mice enable us to analyze the fate of vascular cells during vascular injury^28^. Tracing of the endothelial lineage using the constitutively active systems such as Tie2Cre mice has shown Cre-mediated recombination in endothelial-derived cells and circulating leukocytes^11^. To examine if residential ECs contribute to the neointima formation, we utilized the VE-cadherin (*Cdh5*, also known as CD144)-BAC-Cre^ERT2^-LSL (loxP-stop-loxP)-GFP mouse line (Fig. 2a)^29^. In this system, GFP-labeled residential ECs are only observed in the innermost layer (lumen) without injury, but labeled ECs can be detected in the neointima if they contribute to the neointima formation. Tamoxifen was administered 2 weeks prior to ligation, and the right carotid artery (RCA) and left carotid artery (LCA) were collected at 4 weeks after ligation (Fig. 2b). We first assessed the efficiency of GFP labeling in this model to track residential ECs. Immunostaining for CD31 in the RCA and LCA of the tamoxifen-induced *Cdh5*-BAC-Cre^ERT2^-LSL-GFP mice without ligation revealed that the labeling efficiency, as indicated by GFP-positive ECs, was 42.1% (average for n=3) in CD31^+^ ECs (Fig. 2c-e). After 4 weeks of ligation, GFP-labeled cells were distributed in 34.6% (average for n=4) of the neointima cells in a mottled pattern (Fig. 2f and h). Analysis of serial sections from proximal (close to the ligation point) to distal showed that 29.8% to 44.7% of the neointima cells were GFP-positive in each section (Fig. 2g). Since this frequency was similar to the initial labeling efficiency (Fig. 2e and h), the lineage tracing analysis provides strong evidence that residential ECs contribute to neointima formation after carotid artery ligation.

**Figure 2.**
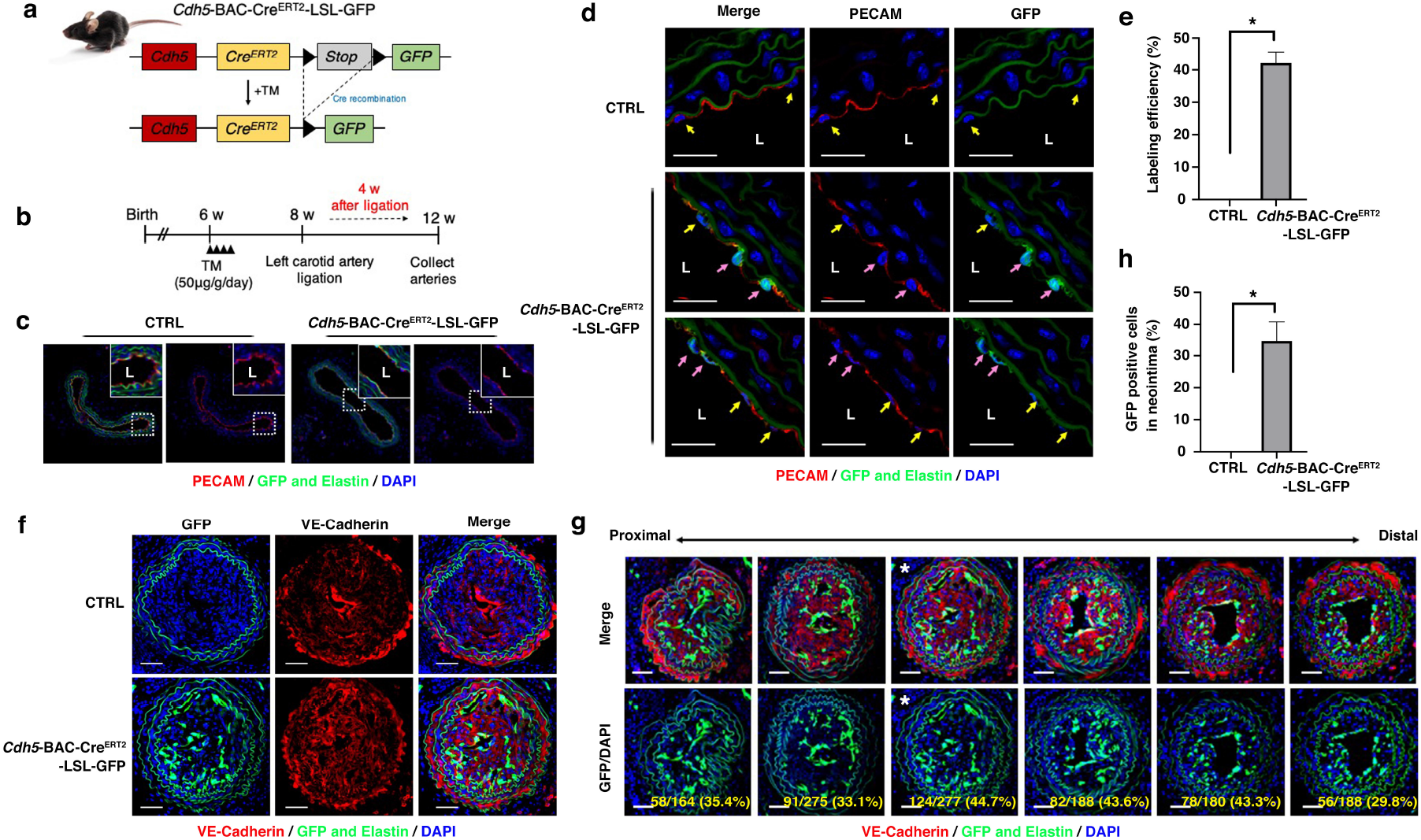
Lineage tracing of ECs during neointima formation upon carotid artery ligation. **(a and b)** Schematic showing the gene-targeting strategy for *Cdh5*-BAC-Cre^ERT2^-LSL-GFP mice and experimental strategy. **(c and d)** Immunostaining of cross sections of un-ligated arteries with PECAM (red) at 6 weeks (n=3) after tamoxifen injection (TM, 50 μg/g ; body weight/day) in RCA (in c) and LCA (in d) of CTRL and *Cdh5*-BAC-Cre^ERT2^-LSL-GFP mice. DAPI (blue), autofluorescence of elastin and GFP (green) are also shown. Pink arrows indicate GFP-positive ECs. Yellow arrows indicate GFP^-^-ECs. **(e)** Quantification of GFP-positive ECs in d. Bar indicates mean ± SEM. \**P*< 0.05, Mann-Whitney test. **(f)** Immunostaining of cross sections of ligated arteries with VE-cadherin (red) at 4 weeks (n=4) after ligation in CTRL and *Cdh5*-BAC-Cre^ERT2^-LSL-GFP mice. DAPI (blue), autofluorescence of elastin and GFP (green) are also shown. **(g)** Serial-sections of LCA in *Cdh5*-BAC-Cre^ERT2^-LSL-GFP mice, which are shown in f. Images are shown in proximal to distal direction from the ligature point. Quantification of GFP-positive cells in neointima using Imaris software is indicated in yellow (number of GFP-positive cells/total number of nuclei). Images with white asterisks are shown in f. All scale bars equal 50 μm. **(h)** Quantification of GFP-positive cells in neointima. Bar indicates mean ± SEM. \**P*< 0.05, Mann-Whitney test.

### CD45 and FSP-1 expression in residential ECs and contribution to neointima formation

ECs display a considerable plasticity to transition to other cell types through various processes, such as EndMT, which induces the loss of certain endothelial markers, such as CD31 and VE-cadherin, and the progressive expression of mesenchymal markers, such as αSMA and FSP-1^11^. Bischoff et al. have reported that CD45 protein PTPase drives EndMT in mitral valve ECs at 6 months after myocardial infarction^18^. Interestingly, CD45-driven EndMT retains an EC phenotype, including CD31 and VE-cadherin expression, and is defined as partial EndMT, which is reversible and can maintain EC functions^20^. To examine the contribution of ECs to neointima formation in more detail, we focused on the earlier stages following carotid artery ligation. One week after ligation, both PECAM and αSMA were expressed in residential ECs of the LCA (Fig. 3a). Since CD45 is not normally expressed in ECs, consistently, we could not detect CD45 in the RCA at 1, 2, or 3 weeks after ligation (Fig. 3b). However, in LCA at 1 week after ligation, CD45 was expressed in approximately 50.3% of the residential ECs (yellow arrowheads in Fig. 3c and d). The deposition of leukocyte-like cells on the endothelium was also observed (blue arrowheads in Fig. 3c), as previously reported^30^, and the developing neointima appeared to be double positive for CD45 and VE-cadherin (yellow arrowheads in Fig. 3e). At 2 weeks after ligation, expression of CD45 and VE-cadherin was observed in the neointima cells, however, the innermost cells were negative for CD45 and VE-cadherin (yellow arrowheads in the inset in Fig. 3f). At 4 weeks after ligation, VE-cadherin expression was confined to the innermost cells, indicating the endothelial lining and re-organization of the lumen (Fig. 3f). The CD45 expression in neointima cells sharply decreased by 4 weeks (Fig. 3f and g). To further examine the expression of EndMT markers in the early stage of neointima formation, we looked at the FSP-1 expression in the RCA and LCA at 1 week after ligation. As expected, FSP-1 was expressed in a monolayer of ECs in the ligated LCA but not in the RCA (Fig. 3h). We further assessed CD45-positive EndMT cells using fluorescence-activated cell sorting (FACS) analysis at 2 weeks after ligation. The population of triple-positive cells (αSMA^+^/CD31^+^/CD45^+^) were significantly increased in the ligated LCAs compared with the contralateral arteries (RCAs) (Fig. 3i and j). These data indicate that CD45-positive EndMT is involved in the initial event in neointima formation upon carotid artery ligation.

**Figure 3.**
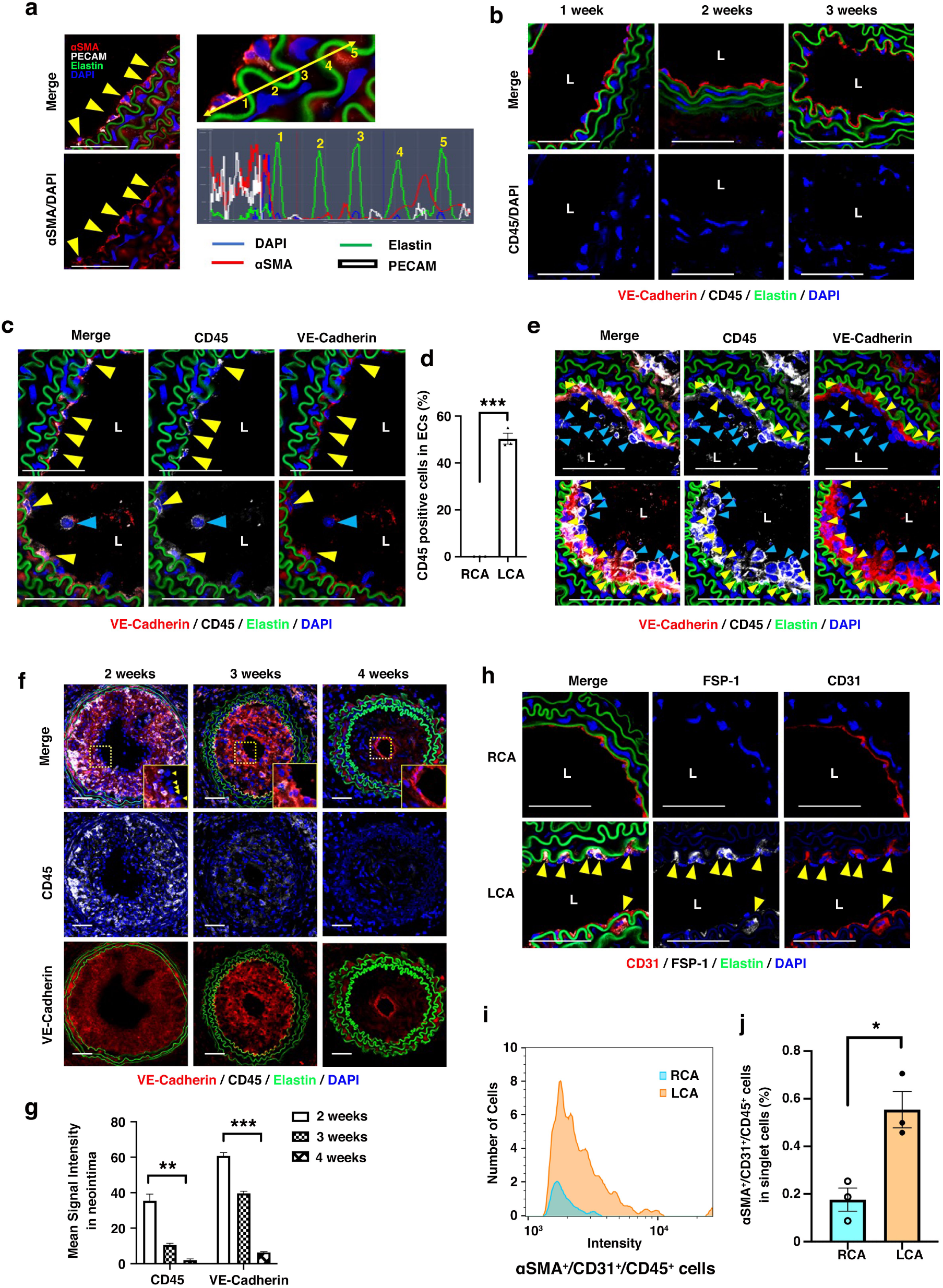
CD45 and FSP-1 are detected in residential ECs and neointima cells upon carotid artery ligation. **(a)** Immunostaining of cross sections of LCA with αSMA (red) and PECAM (white) at 1 week (n=5) after ligation in WT mice. Histogram shows signal intensity on the yellow line. **(b)** Immunostaining of cross sections of RCA with VE-cadherin (red) and CD45 (white) at 1 week (n=5), 2 weeks (n=5), and 3 weeks (n=4) after ligation in LCA of WT mice. **(c, e, and f)** Immunostaining of cross sections of ligated arteries with VE-cadherin (red) and CD45 (white) at 1 week (n=5, in c and e), 2 weeks (n=5, in f), 3 weeks (n=4, in f), and 4 weeks (n=4, in f) after ligation in LCA of WT mice. Yellow arrowheads (in c and e) indicate VE-Cadherin^+^/CD45^+^ cells. Blue arrowheads (in c and e) indicate VE-cadherin^-^/CD45^+^ cells. Yellow arrowheads (in f) indicate VE-cadherin^-^/CD45^-^ cells in the innermost layer at 2 weeks after ligation. **(d)** Quantification of CD45-positive cells in ECs is shown in b and c, \*\*\**P*< 0.001, unpaired *t*-test. **(g)** Quantification of mean signal intensity for CD45 and VE-cadherin are shown in f. \*\**P*< 0.01, \*\*\**P*< 0.001, two-way ANOVA. **(h)** Immunostaining of cross sections of ligated arteries with CD31 (red) and FSP-1 (white) at 1 weeks (n=5) after ligation in WT mice. Yellow arrowheads indicate CD31^+^/FSP-1^+^ cells. **(a-h)** All scale bars equal 50 μm. DAPI (blue) and autofluorescence of elastin (green) are also shown. **(i)** Representative graph of flow cytometry (FACS) of αSMA^+^/CD31^+^/CD45^+^ cells in RCA and LCA at 2 weeks (n=3) after ligation in WT mice. **(j)** Histogram shows αSMA^+^/CD31^+^/CD45^+^ cells in RCA and LCA of total singlet cells. Bar indicates mean ± SEM. \**P*< 0.05, unpaired *t*-test.

### EndMT markers and pSmad activation in the early stage of neointima formation

To examine the expression of other EndMT markers during neointima formation, we performed quantitative PCR (qPCR) analysis. RNA was extracted from the RCA and LCA at 2 weeks after ligation or without ligation as a control (CTRL). The expression of SMC contractile genes was low in the LCAs from the mice with ligation, whereas the expression of *Cnn1* and *Myh11* were increased in the contralateral arteries (RCAs) (Fig. 4a and Extended Data Fig. 2). Transforming growth factor-β (TGFβ) has been linked to the induction of EndMT; among the TGFβ family, *Tgfβ1* and *Tgfβ3* were significantly increased in the LCA from the ligated mice, with *Tgfβ1* showing a 15-fold change compared with that of the CTRL and contralateral artery (RCA). The expression of *Snail* (also known as Snail1) and *Slug* (also known as Snail2) were higher in the LCAs from the ligated mice than the control mice, and mRNA expression of *Fsp1, Thbs1*, and *CD45* was also high in the LCAs from these mice (Fig. 4a and Extended Data Fig. 2). Consistent with previous reports, the protein levels of Thbs1 and phosphorylation (p) of extracellular signal-regulated kinase (ERK) were increased in the ligated LCA, and expression of YAP was induced after ligation (Fig. 4b)^5, 26, 31^. In addition, we found that the protein levels of Slug and pSmad were increased in the ligated LCA (Fig. 4b). To further elucidate the activation of the TGFβ signaling pathway, we examined the pSmad2/3 levels in neointima cells. In inner cells at 1 week after ligation, nuclear pSmad2/3 (active pSmad2/3) was detected in the LCA, whereas it was not detected in the contralateral artery (RCA) (Fig. 4c). Interestingly, some inner cells of LCA were CD45-positive but retained pSmad2/3 in the cytoplasm (yellow arrows in Fig. 4c). Furthermore, as shown in Fig. 4d, pSmad2/3 was not activated in medial SMCs, but pSmad2/3 was present in the nuclei of neointima cells in the tamoxifen-induced *Cdh5*-BAC-Cre^ERT2^-LSL-GFP mice at 4 weeks after ligation. These results strongly suggest that increases in TGFβ signaling and CD45-positive EndMT contribute to the initiation of neointima formation.

**Figure 4.**
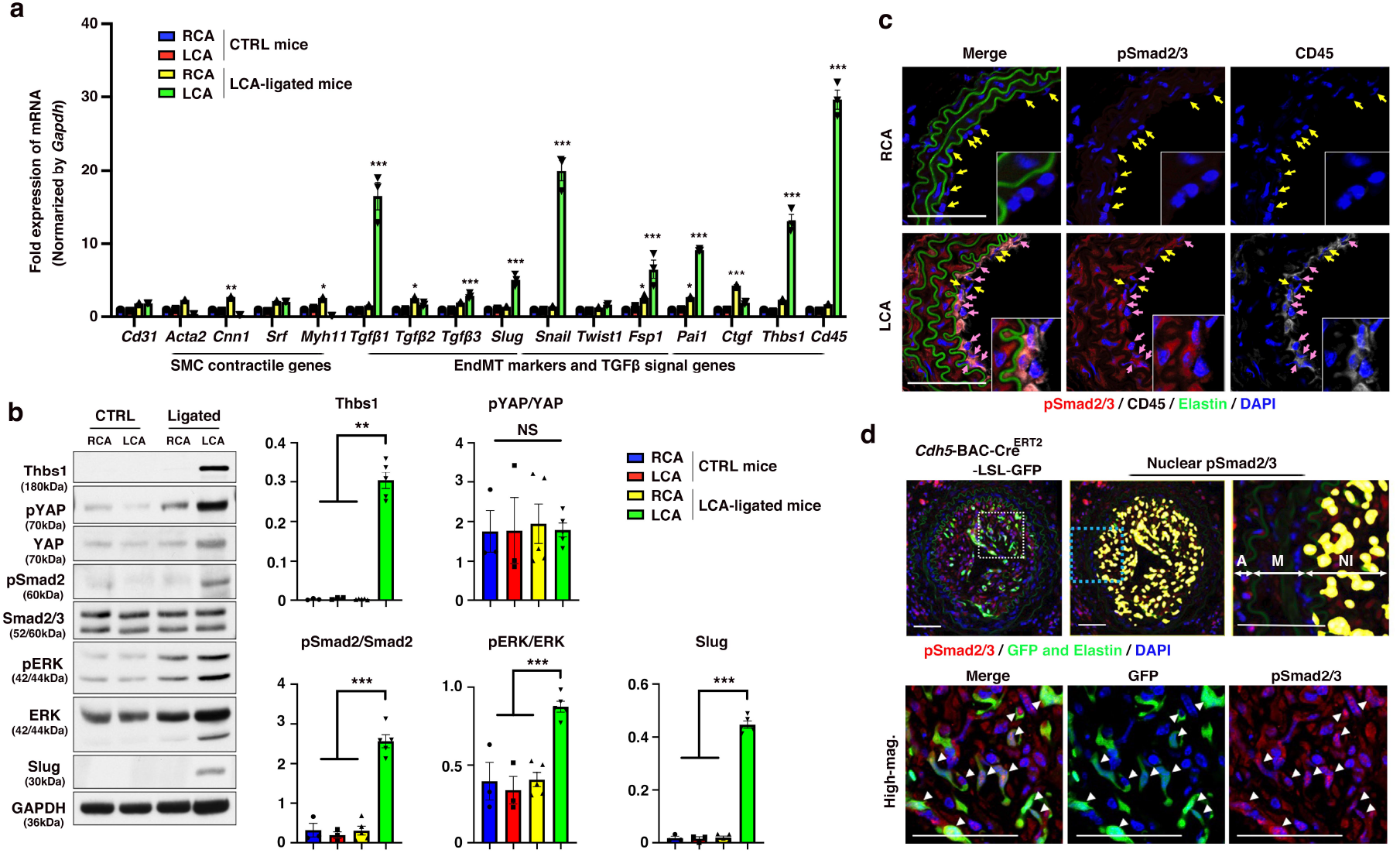
EndMT marker genes and nuclear pSmad are prominent in developing neointima. **(a)** Quantitative polymerase chain reaction (qPCR) analysis of CD31, SMC contractile genes, EndMT markers, and TGFβ signal genes from right carotid artery (RCA) or left carotid artery (LCA) of CTRL (n=3) and LCA-ligated mice (n=3). Bars indicate mean ± SEM. \*\*\**P*< 0.001, \*\**P*< 0.01, two-way ANOVA. Individual graphs are shown in Extended Data Fig. 2. **(b)** Representative western blots show Thbs1, pYAP, pERK pSmad2, and Slug levels from RCA or LCA in CTRL (n=3) and LCA-ligated mice (n=5). Quantification graph is shown on the right. Bars indicate mean ± SEM. \*\*\**P*< 0.001, \*\**P*< 0.01, one-way ANOVA and the Kruskal-Wallis test for Thbs1. **(c)** Immunostaining of cross sections of ligated arteries with pSmad2/3 (red) and CD45 (white) at 1 week (n=5) after ligation in WT mice. DAPI (blue) and autofluorescence of elastin (green) are also shown. Yellow arrows show EC without nuclear pSmad2/3 and pink arrows indicate EC with nuclear pSmad2/3. **(d)** Immunostaining of cross sections of ligated arteries with pSmad (red) at 4 weeks (n=4) after ligation in *Cdh5*-BAC-Cre^ERT2^-LSL-GFP mice. DAPI (blue), autofluorescence of elastin and GFP (green) are also shown. Highly magnified images in the dashed white box are shown at the bottom. White arrow heads indicate GFP-positive cells with nuclear pSmad2/3. On the right, the cells with nuclear pSmad2/3 were detected by using Imaris software and are indicated in yellow. Highly magnified image in dashed light-blue box is shown on the right. NI: neointima, M: medial layer, A: adventitial layer. All scale bars equal 50 μm.

### Stabilization of hypoxia-inducible factor-1 alpha (HIF-1α) induces expression of CD45 during EndMT in human aortic ECs

We next investigated the stimuli that trigger CD45-positive EndMT using human aortic ECs (HAECs). Recent studies have indicated that matrix stiffness alters the phenotype of ECs and promotes EndMT^32, 33^. Therefore, we first cultured HAECs on hydrogels with varying levels of cell matrix stiffness. HAECs cultured on soft gels (2 kPa) showed a spherical morphology and higher expression of EndMT markers such as *TGFβ1, TGFβ3, SNAIL*, and plasminogen activated inhibitor-1 (*PAI1*) than HAECs cultured on hard gels (120 kPa and 30 kPa); however, *CD45* expression was not altered on soft or hard gels (Extended Data Fig. 3). Next, we examined if hypoxic conditions induce CD45-positive EndMT. HAECs were treated with 100 μM cobalt chloride (CoCl_2_), a HIF activator^34^, or 100 nM of another HIF activator^35^, VH298, for 48 h, and an increase in the expression of EndMT markers, including *TGFβ1, TGFβ3, SNAIL*, and *PAI1*, and upregulation of CD45 expression were observed (Fig. 5a and Extended Data Fig. 4). Similar results were obtained with human cardiac artery EC (HCAEC) and human pulmonary artery EC (HPAEC) (Extended Data Fig. 5). Although Bischoff et al. previously reported that mitral valve ECs, but not arterial ECs, develop CD45-positive EndMT after 4 days in culture with 1 ng/ml TGFβ1^18^, the expression of EndMT markers and CD45 was not upregulated in HAECs by treatment with 1 ng/ml recombinant human (rh) TGFβ1 or 180 ng/ml rhTHBS1 for 48 h (Fig. 5a). Treatment with 100 μM CoCl_2_ or 100 nM VH298 for 48 h increased the protein expression of HIF-1α, THBS1, SLUG and phosphorylation of vascular endothelial growth factor receptor 2 (pVEGFR2), but the expression level of VE-cadherin was not changed (Fig. 5b). These results suggest that hypoxic conditions with CoCl_2_ or VH298 which mimicked ligated artery, promotes stabilization of HIF-1α and triggers CD45-positive EndMT. Consistent with our observation, the protein levels of HIF-1α increased in the ligated LCA but were not detected in the control carotid arteries (without ligation) or the contralateral arteries (RCAs) at 2 weeks after ligation (Fig. 5c). Additionally, the levels of fibronectin, collagen type I, and the endoplasmic reticulum (ER)-stress marker KDEL (BiP; immunoglobulin heavy chain-binding protein), which have been associated with the induction of EndMT^36, 37^, were increased in the ligated LCAs (Fig. 5c). Immunostaining in the early or developed stages of neointima formation showed that HIF-1α was strongly expressed in the EC monolayer at 1 week after ligation, and in the innermost cells of the established neointima at 4 weeks after ligation (Fig. 5d). Taken together, these results indicate that stabilization of HIF-1α by hypoxia associated with blood flow cessation may induce CD45-positive EndMT and trigger neointima formation upon carotid artery ligation.

**Figure 5.**
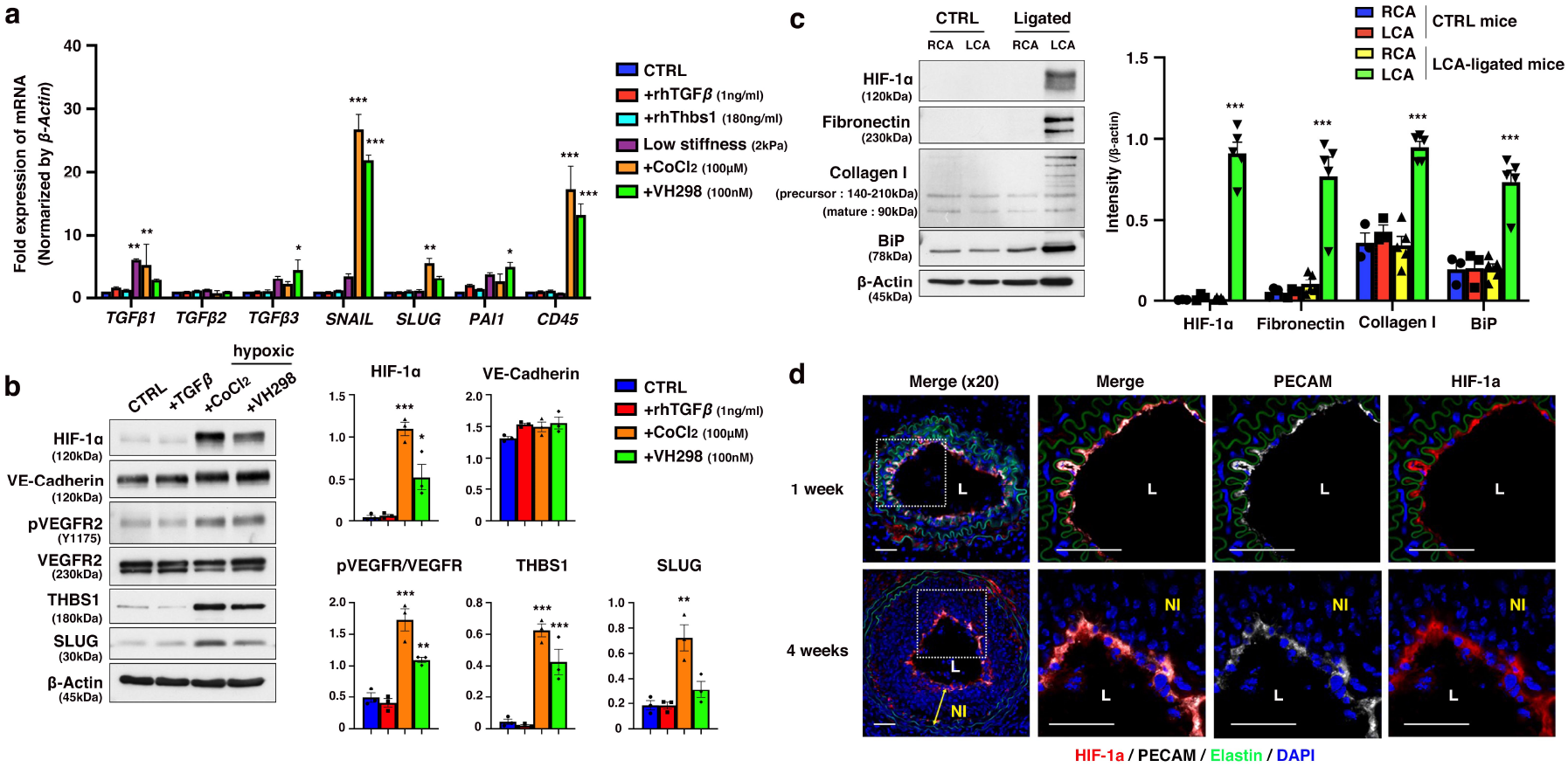
Hypoxic conditions induce EndMT markers with CD45 expression in human aortic endothelial cells (HAECs) and ligated carotid artery. **(a)** qPCR analysis of EndMT markers, TGFβ signal genes, and CD45. HAECs were cultured with recombinant human TGFβ1 (rhTGFβ, 1 ng/ml) or rhTHBS1 (180 ng/ml) or CoCl_2_ (100 μM) or von Hippel-Lindau Tumor Suppressor (VHL) inhibitor (VH298, 100 nM) for 48 h or cultured on soft hydrogel (2 kPa) for 24 h. n=3 for each condition. Bars indicate mean ± SEM. \*\*\**P*< 0.001, two-way ANOVA. **(b)** HAECs were cultured with rhTGFβ (1 ng/ml) or CoCl_2_ (100 μM) or VH298 (100 nM) for 48 h. Representative western blot shows HIF-1α, VE-cadherin, pVEGFR2, VEGFR2, THBS1, and SLUG levels in CTRL or TGFβ1 or hypoxic conditions (n=3 for each condition). Quantification graph is shown on the right. Bars indicate mean ± SEM. \*\*\**P*< 0.001, \*\**P*< 0.01, one-way ANOVA. **(c)** Representative western blot shows HIF-1α, Fibronectin, Collagen, and KDEL (BiP) levels from RCA or LCA in CTRL (n=3) and LCA-ligated mice (n=5). Quantification graph is shown on the right. Bars indicate mean ± SEM. \*\*\**P*< 0.001, one-way ANOVA. **(d)** Immunostaining of cross sections of ligated arteries with HIF-1α (red) and PECAM (white) at 1 week (n=5) and 4 weeks (n=4) after ligation in WT mice. DAPI (blue) and autofluorescence of elastin (green) are also shown. Highly magnified images in the dashed white box are shown on the right. Yellow arrow indicates neointima (NI) layer. L: lumen. All scale bars equal 50 μm.

### PTPase activity of CD45 promotes integrin α11-SHARPIN complex formation and regulates VE-cadherin stability

Integrins act as biomechanical sensors of the microenvironment and function as a structural and biochemical bridge between the ECM and cytoskeletal linker proteins^2^. To investigate the changes in integrins under hypoxic condition, we examined the expression levels of integrin subunits in HAECs under CTRL and 100 μM of CoCl_2_ treatment after 48 hours of culture by qPCR. The results showed that the expression levels of the integrins α1, α11, αIIb, and β6 were significantly upregulated by CoCl_2_ treatment (Fig. 6a). As shown in Fig. 6b, integrins α1 and α11 are known as collagen receptors and form dimers with integrin β1 (α1β1, α11β1)^38^. Integrin αIIb forms a dimer with integrin β3 (αIIbβ3) and is known as a fibrinogen receptor that promotes blood clot formation upon binding^39^. Integrins β6 and β8 form dimers with integrin αv (αvβ6, αvβ8) and are known to play a role in converting inactive (latent) TGFβ to the active form^40^. We next examined the relationship between these integrins and CD45 by using an CD45 PTPase inhibitor. CoCl_2_-induced integrin α11 expression was completely abolished by this inhibitor (30 nM, Fig. 6c), suggesting that integrin α11 upregulation by CoCl_2_ treatment is dependent on the PTPase activity of CD45. In addition, we examined the effect of inhibition of CD45 PTPase activity or TGFβ activation on the expression of EndMT markers under CoCl_2_ treatment. Treatment with a CD45 PTPase inhibitor did not alter the expression of EndMT markers, but treatment with SB525334, a TGFβ/Smad inhibitor, significantly suppressed the expression of CoCl_2_-induced EndMT markers (Fig. 6d). The upregulation of CD45 expression by CoCl_2_ was also suppressed by SB525334 treatment, suggesting that the TGFβ/Smad pathway regulates CD45 expression under CoCl_2_ treatment.

**Figure 6.**
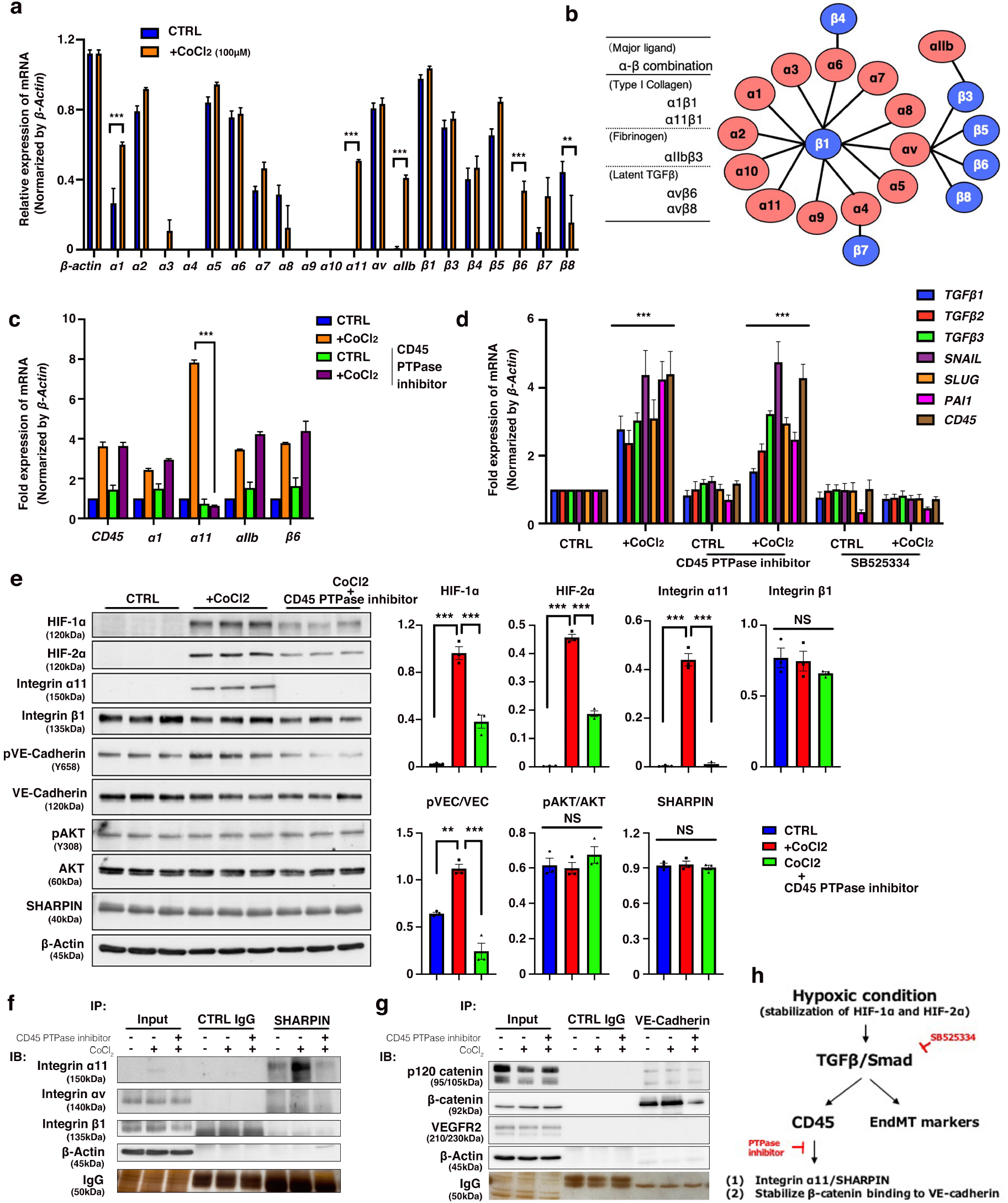
CD45 PTPase activity promotes integrin α11-SHARPIN complex formation and regulates VE-cadherin stability. **(a)** Relative mRNA expression (β-actin as a 1.0) of integrin subunits for CTRL or CoCl_2_ (100 μM) conditions in HAECs (n=5). Bars indicate mean ± SEM. \*\*\**P*< 0.001, \*\**P*< 0.01, two-way ANOVA. **(b)** Possible heterodimeric integrin subtypes and diagrammatic representation of a known integrin heterodimer. **(c)** qPCR analysis of integrin α1, α11, αIIb, β6, and CD45 with or without CD45 PTPase inhibitor (30 nM) under CoCl_2_ treatment for 48 h. n=3 for each condition. Bars indicate mean ± SEM. \*\*\**P*< 0.001, two-way ANOVA. **(d)** qPCR analysis of EndMT markers, TGFβ signal genes, and CD45. HAECs were cultured with CoCl_2_ (100 μM) for 48 h, and with or without CD45 PTPase inhibitor (30 nM) or SB525334 (10 nM). n=3 for each condition. Bars indicate mean ± SEM. \*\*\**P*< 0.001, two-way ANOVA. **(e)** Western blots show HIF-1α, HIF-2α, integrin α11, β1, VE-Cadherin, AKT, and SHARPIN levels in CTRL, CoCl_2_ (100 μM), and CoCl_2_ with CD45 PTPase inhibitor (30 nM). n=3 for each condition. Quantification graph is shown on the right. Bars indicate mean ± SEM. \*\*\**P*< 0.001, one-way ANOVA and Kruskal-Wallis test for HIF-2α and pVEC/VEC. **(f-g)** Immunoprecipitation (IP) with anti-SHARPIN (in f), anti-VE-cadherin (in g), or control immunoglobulin G (IgG) followed by western blotting for integrin α11, αv, and β1 (in f), and p120 catenin, β-catenin, and VEGFR2 (in g). n=2 for each IP. Silver staining shows the heavy chain of IgG in IP lysates. IB: immunoblot. **(h)** Schematic diagrams of hypoxic condition-induced partial EndMT.

To investigate the effect of the CD45 PTPase inhibitor on intracellular signaling pathways, we performed western blot analysis. As shown in Fig. 6e, integrin α11 expression was markedly upregulated under CoCl_2_ treatment and was completely suppressed by CD45 PTPase inhibitor, consistent with the results of qPCR analysis. The phosphorylation of Y658, which contributes to the stabilization of VE-cadherin, was enhanced by CoCl_2_ but inhibited by CD45 PTPase inhibitor. Interestingly, the expression of HIF-1α and HIF-2α, which are stabilized by CoCl_2_ treatment, was significantly suppressed by CD45 PTPase inhibitor. On the other hand, no changes were observed in the expression of integrin β1 and Shank associated RH domain-interacting protein (SHARPIN, known as a negative regulator of integrin β1) or the phosphorylation levels of AKT (Fig. 6e). Similarly, the phosphorylation of phosphoinositide 3-kinase (PI3K), Src, phosphatase and tensin homolog (PTEN) and VEGFR2 and the expression level of kruppel-like factor 4 (KLF4) were not altered by CD45 PTPase inhibitor (Extended Data Fig. 6). To further investigate the role of CD45 in the induction of integrin α11 expression under CoCl_2_ treatment, we performed immunoprecipitation (IP) using a SHARPIN antibody. Recent work from Lerche et al. shows that SHARPIN binds to integrin α11, thereby regulating the mechanosensory function of mammary gland fibroblasts^41^. We found that SHARPIN binds to integrin α11 under CoCl_2_ treatment, but not to integrin αv or β1, and SHARPIN binding to integrin α11 was suppressed by treatment with a CD45 PTPase inhibitor (Fig. 6f). This result suggests that CD45 PTPase activity facilitates integrin α11-SHARPIN complex formation and may contribute to partial EndMT under CoCl_2_ treatment. To examine the EC stability in CoCl_2_ treatment, we performed IP using a VE-cadherin antibody. While β-catenin binding to VE-cadherin was maintained under CoCl_2_ treatment, β-catenin was released from VE-cadherin by inhibition of CD45 PTPase activity (Fig. 6g). Collectively, these findings suggest that stabilization of HIF-1α (and/or HIF-2α) by CoCl_2_ treatment induces integrin α11 expression in a CD45 PTPase activity-dependent manner and promotes its binding to SHARPIN. β-catenin also maintains the complex formation with VE-cadherin in this condition (termed partial EndMT). In contrast, under the inhibition of CD45 PTPase, β-catenin is released from VE-cadherin and weakens cell-cell junctions (termed complete EndMT) (Fig. 6h). These data indicate that CD45 PTPase activity promotes integrin α11-SHARPIN complex formation and regulates cell-cell junction stability undergoing EndMT process.

### Partial EndMT sustains cell-cell junctions and ability of tube formation

To investigate the cell morphology and cell-cell junctions of HAECs in CoCl_2_ treatment with or without CD45 activity, we performed immunostaining for VE-cadherin and phalloidin. The CTRL cells showed strong expression of VE-cadherin and a rhombic morphology with cortical actin distribution, whereas HAECs were elongated and had expanded cell areas with moderate expression of VE-cadherin and moderate actin fibers under CoCl_2_ treatment (partial EndMT, Fig. 7a-c). Inhibition of CD45 PTPase activity resulted in a similar elongated morphology to CoCl_2_ treatment alone (Fig. 7b), however, the cells showed a significant decrease in VE-cadherin expression and prominent actin stress fibers (complete EndMT, Fig. 7a and c). To support this observation, we assessed the mechanical properties of cells undergoing partial EndMT (in HAECs) using atomic force microscopy. The height of the cells undergoing partial EndMT was lower than that of the other cells (Fig. 7d, e). The Young’s modulus (stiffness) of these cells was increased compared with that of the CTRL cells but was comparable to that of the cells with complete EndMT (Fig. 7d, f). When cell-substrate adhesion is preferentially maintained, cell height negatively correlates with cell tension^42^, therefore we analyzed the correlation between height and stiffness. Interestingly, the cells in the CTRL and complete EndMT groups were negatively correlated; however, the cells with partial EndMT did not show any correlation between the height and tension (Fig. 7g). This result indicated that sustained cell-cell junctions in cells undergoing partial EndMT disrupted the balance between cell-cell junctions and cell-substrate adhesions, thereby abolishing the negative correlation. Finally, we functionally assessed whether partial EndMT retains ECs function. HAECs were treated with 100 μM CoCl_2_ to induce EndMT either with or without 30 nM CD45 PTPase inhibitor. Forty-eight hours later, HAECs were seeded onto Matrigel-coated 24-well plates, and tube formation was evaluated after 8 hours (Fig. 7h). While tube formation occurred in the CTRL and partial EndMT groups, the cells with complete EndMT lost their ability to form tubes (Fig. 7h, i). These results indicate that partial EndMT enables cells to maintain cell-cell junctions during vascular remodeling.

**Figure 7.**
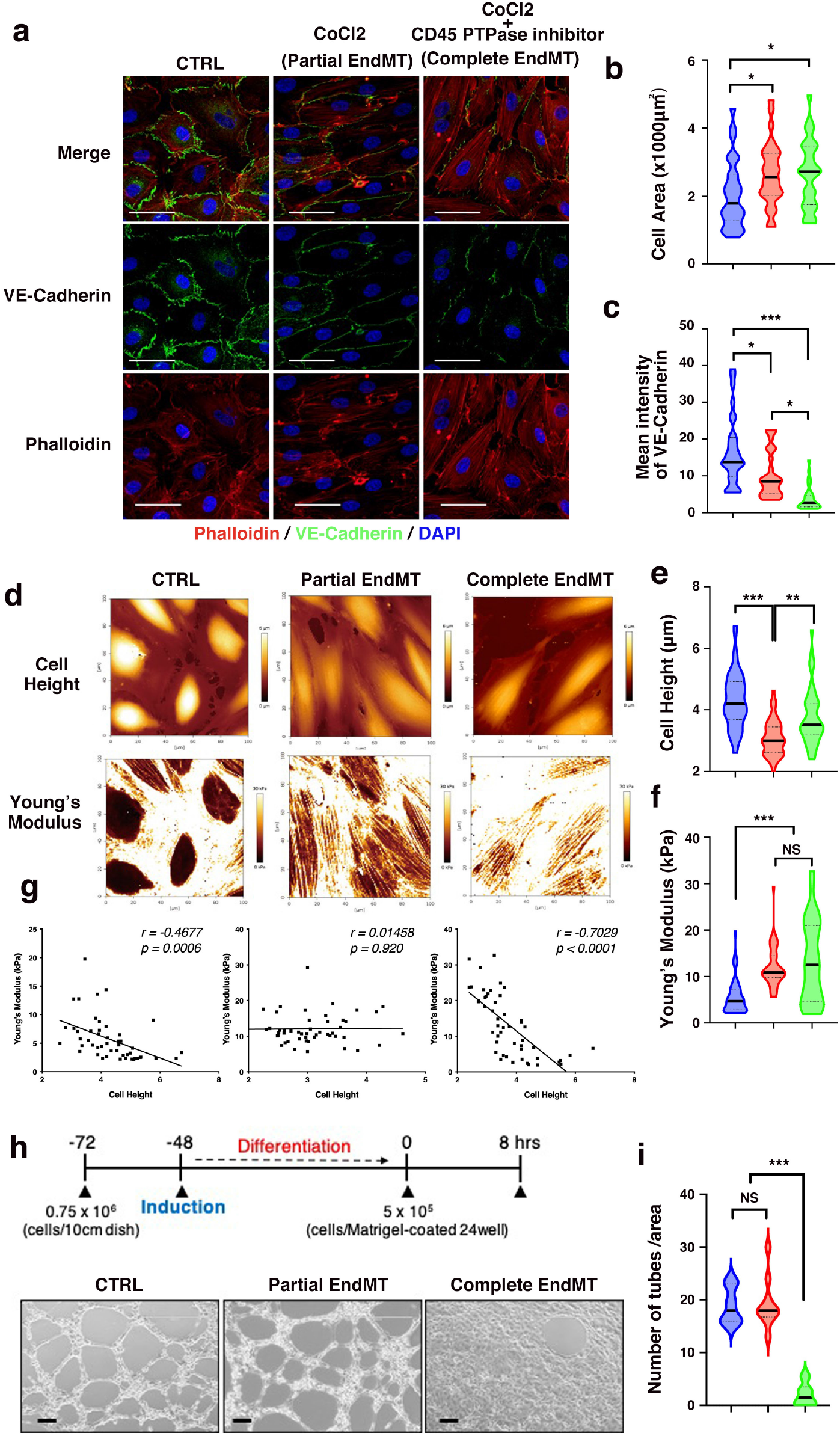
Partial EndMT sustains endothelial functions. **(a)** Representative immunostaining with phalloidin (red), VE-cadherin (green), and DAPI (blue). Scale bars equal 50 μm. **(b and c)** Violin plots for quantification of cell area and mean intensity of VE-cadherin are shown on the right. CTRL (blue), CoCl_2_ (partial EndMT: red) and CoCl_2_ with 30 nM of CD45 PTPase inhibitor (complete EndMT: green). 30 cells and cell-cell junctions were randomly selected for quantification. Median is indicated by bold black bar. \*\*\**P*< 0.001, \**P*< 0.05, one-way ANOVA for b and Kruskal-Wallis test for c. **(d)** Young’s modulus of actin fibers in CTRL, CoCl_2_ (partial EndMT), or CoCl_2_ with CD45 PTPase inhibitor-treated cells (complete EndMT) (n=50 per conditions) measured using atomic force microscopy. Representative topographic images and stiffness maps are shown. (**e and f**) Violin plots for quantification of cell height and Young’s modulus are shown on the right. CTRL (blue), partial EndMT (red) and complete EndMT (green). Median is indicated by bold black bar. \*\*\**P*< 0.001, \*\**P*< 0.01, Kruskal-Wallis test. NS: not significant. (**g**) Graphs indicating correlation with cell height and Young’s modulus are shown at the bottom. Pearson *r* shows correlation, where +1 is total positive linear correlation, 0 is no linear correlation, and -1 is total negative correlation. Two-tailed *P* values are shown in graph. **(h)** Matrigel-based tube formation assay. Representative images of CTRL, CoCl_2_ (partial EndMT), or CoCl_2_ with CD45 PTPase inhibitor-treated cells (complete EndMT) (n=10 per conditions) that were plated on Matrigels for 8 h. (**i**) Violin plots for quantification of the number of tubes are shown on the right. CTRL (blue), partial EndMT (red) and complete EndMT (green). Median is indicated bold black bar. \*\*\**P*< 0.001, one-way ANOVA. NS: not significant.

## Discussion

In this study, we used EC-lineage tracing system and revealed that residential ECs contribute to neointima formation after carotid artery ligation. In the early stage of neointima formation, hematopoietic marker CD45 was detected in cells undergoing EndMT and activation of TGFβ signaling was prominent in the neointima cells. In vitro system using primary human ECs, CD45-positive EndMT was induced by stabilization of HIF-1α with CoCl_2_ or VH298, which mimicked hypoxic condition of ligated artery, and promoted integrin α11 expression and complex formation with SHARPIN. Pharmacological inhibition of CD45 PTPase activity did not affect the expression of EndMT markers but 1) suppressed expression of integrin α11 and abolished complex formation with SHARPIN, and 2) disrupted the binding between β-catenin and VE-cadherin, thereby destabilizing cell-cell junctions and promoted complete EndMT. Our study proposes a novel regulatory mechanism of partial EndMT and provides evidence that CD45-positive EndMT plays a crucial role in lumen re-organization associated with pathological vascular remodeling (Fig. 8).

**Figure 8.**
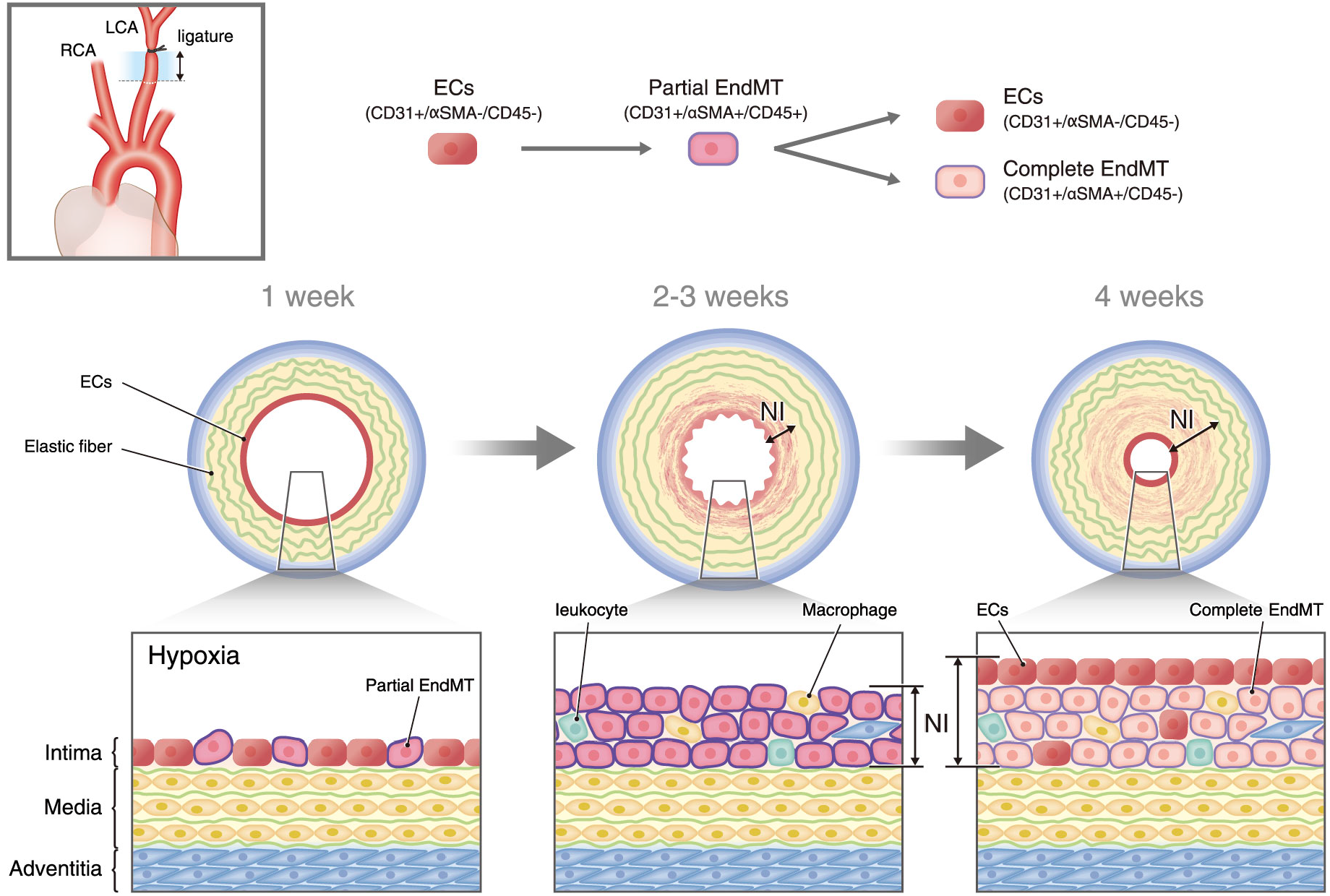
A model illustrating the contribution of partial EndMT in lumen re-organization after carotid artery ligation. Hypoxic condition (stabilization of HIFs) induces CD45-positive EndMT (partial EndMT) at 1 week after ligation. At 2-3 weeks after ligation, ECs undergoing partial EndMT proliferate and form neointima (NI). At 4 weeks after ligation, CD45 expression on ECs is rapidly down-regulated and ECs complete EndMT. A new lumen is re-established by ECs.

Carotid artery stenosis is known to reduce blood flow to the brain and cause strokes. Surgical treatments include carotid endarterectomy and carotid artery stenting, both of which aim to open up the stenotic area^43^. Alternatively, prevention of risk factors for atherosclerosis, such as hypertension, hyperlipidemia (dyslipidemia), and diabetes, as well as the use of antiplatelet agents, is a common internal medical treatment strategy for carotid artery stenosis. Since the origin of neointima cells during carotid artery stenosis has long been debated, accurate elucidation of the pathogenesis of this disease is needed to develop treatment methods and achieve early detection of stenosis. Initially, mice deficient in genes such as matrix metalloproteinase (*Mmp*)*-2, Mmp-9*, plasminogen activator inhibitor (*Pai*)*-1*, and vitronectin (*Vtn*) showed inhibition of neointima formation^44-46^, and it was actively discussed that SMCs in the tunica media migrated to the luminal side, accompanied by αSMA expression. Subsequently, mice deficient in vascular SMC-specific (smCre-Tg) Notch1 were generated and evaluated for neointima formation, but neointima formed despite some inhibition^47^. Therefore, it necessitated the use of a cell-type specific lineage tracing system or gene-deletion system to precisely identify the origin of neointima cells and factors involved in the neointima formation. Furthermore, indirect contribution of ECs has been reported as the mechanism of neointima formation. Janmaat et al. showed that a combination of erythropoietin (EPO) and inflammatory cytokine, TNF-α, induces a release of PDGF-BB from ECs, which in turn phosphorylates Stat5 (Signal transducer and activator of transcription 5) in SMCs and promotes SMC proliferation^48^. Anggrahini et al. reported that EC-derived endothelin-1 (ET-1) promoted SMC proliferation during neointima formation^49^. Interestingly, they observed that the innermost layer of cells became CD45 positive early in the stenosis (3 days after ligation), and they discussed the possibility of leukocyte deposition as a trigger of ET-1 production on ECs. Kumar et al. reported that the neointima cells were CD45 positive, but they determined that this phenomenon resulted from leukocyte infiltration^50^. Noma et al. also reported that leukocyte deposit and infiltrate the intima in the early stages (3 days after ligation) of neointima formation^27^; however, the possibility of EndMT had not been addressed. While Yoshida et al. first proposed a solo role of ECs in the regulation of neointima formation by demonstrating that EC-specific (Tie2-Cre) deletion of *Klf4* attenuated neointima formation^51^, they did not identify the origin of neointima cells. In this study, we provided evidence that ECs directly contribute to the neointima formation by applying a GFP-labeling (tracing) system for ECs (Fig. 2f). Although we observed deposition of leukocytes on residential ECs as previously reported, we showed that monolayer ECs were positive for CD45 without deposition of leukocytes and CD45-positive EndMT resulted in neointima formation (Fig. 3c).

CD45 is a 180-220 kDa of a single-transmembrane protein with O-linked and N-linked glycosylation sites on the extracellular domain and two phosphotyrosine PTPase sites on the intracellular domain (Extended Fig. 4a)^52^. The functional difference in CD45 isoforms, which are due to alternative splicing of the extracellular domain, is not well investigated in the current studies. CD45 is known to regulate cell growth, migration, and differentiation by modulating downstream Src kinases through its PTPase activity in hematopoietic cells, including macrophages^53^. We also examined the effect of Src phosphorylation and its downstream kinase under CoCl_2_ treatment in HAECs by blocking CD45 PTPase activity, but no differences were detected between CoCl_2_ treatment with or without the CD45 PTPase inhibitor (Extended Fig. 6). A more detailed examination of initial changes and changes over time may be necessary to elucidate the downstream events of CD45 under CoCl_2_ treatment. Proximity-dependent biotin identification (BioID or TurboID and related tools) for CD45 would be useful for further characterization of interacting molecules.

In general, CD45 is expressed on leukocytes, including lymphocytes, eosinophils, monocytes, basophils, and neutrophils. Although CD45 is not expressed on ECs, it is known to be expressed on hemangioblasts in the yolk sac, placenta, and dorsal aorta during embryonic stages^54, 55^. In adult tissues, CD45 was expressed on sheep mitral valve ECs at 6 months after myocardial infarction, as shown by Bischoff et al.^18^ In their experiments, CD45-positive EndMT was induced by treatment with 1 ng/ml TGFβ1 for 4 days, and TGFβ1-induced EndMT was completely suppressed by 1 µM CD45 PTPase inhibitor treatment. However, in our present study using HAECs, CoCl_2_-induced EndMT was not suppressed by 30 nM of CD45 PTPase inhibitor (Fig. 6d). This discrepancy may be due to the inhibitor concentration (1 µM vs 30 nM) or differences in response by cell type (mitral valve ECs vs HAECs) or other conditions (TGFβ1 for 4 days vs CoCl_2_ for 2 days). Pi et al. reported that EC-specific connective tissue growth factor (CTGF) is required for vascular remodeling associated with PAH in mice^56^. They generated *Cdh5*-Cre-*Ctgf*^flox/flox^ mice and *Ctgf*-GFP mice and showed that EC-specific knockout of *Ctgf* protected against PAH and chronic hypoxia induced PAH. They also showed that some of the CD45-positive lung cells are GFP (CTGF) and CD11b-positive, indicating that ECs may not be the only CTGF-expressing cells and concluded that CTGF contributes to the expansion of hematopoietic cell pool within the lung. However, the possibility of CD45-positive EndMT triggered the development of PAH cannot be ruled out in their model. Since CD45 is routinely used as a negative marker for ECs, existence of CD45-positive ECs may have been overlooked in the previous studies.

Integrin α11, which is not expressed in normal ECs but is markedly upregulated by CoCl_2_ treatment, is a known marker of cancer-associated fibroblasts (CAFs) as a collagen-binding mesenchymal integrin. It has been reported to be involved in myofibroblast differentiation, matrix reorganization, collagen deposition, and fibrogenesis^57-59^. Recently, integrin α11 has been shown to form a complex with the tyrosine kinase receptor PDGFRβ, regulates c-Jun N-terminal kinase (JNK) signaling and tenascin-C production, and promotes breast cancer progression^60^. In the present study, we showed that the expression of integrin α11 is regulated in a CD45 PTPase activity-dependent manner (Fig. 6c and e). We speculate that upregulation of integrin α11 induces binding to cell surface receptor(s) and alters ECM production/distribution and intracellular signal(s), and allows EndMT to proceed while maintaining cell-cell junctions. However, the detailed mechanism(s), such as identification of the CD45-substrate and how it regulates integrin α11 expression, is still unclear. More recently, Lerche et al. reported that the expression of integrin α11 is regulated by the presence of SHARPIN and controls cell spreading and stiffness^41^. SHARPIN is an adapter protein known as an integrin inhibitor that binds specifically to the α-tail (GFFKR amino acid sequence) and inhibits the recruitment of talin and kindlin to the β-tail of integrin β1^61, 62^, thereby prohibiting maturation of focal adhesions. The sustained β-catenin binding to VE-cadherin observed under CoCl_2_ treatment is dependent on the CD45 phosphatase activity. These findings suggest that the presence of integrin α11/SHARPIN complex inhibits activation of integrin β1 and locally prohibits stress fiber formation, which may counteract the completion of the EndMT process. Therefore, more follow-up studies are necessary to fully determine how CD45 regulates integrin α11 expression and how integrin α11/SHARPIN controls β-catenin binding to VE-cadherin.

## Methods

### Mice

C57BL/6J (wild-type) mice were purchased from Charles River, Japan. VE-cadherin (*Cdh5*)-BAC-Cre^ERT2^ mice were generated previously^29^ and crossed with *flox*-stop-*flox* (LSL)-GFP mice^63^. For lineage tracing study, 100 μl of tamoxifen (Sigma-Aldrich, T5648) dissolved in oil at 10 mg/ml was injected (final concentration ; 50 μg/g body weight) into the inguinal fat for 4 consecutive days, 2-weeks prior to carotid artery ligation. Male mice were used for carotid artery ligation in the Fig. 1, Fig. 3 and Fig. 4 to exclude hormonal regulation by estrogen. Male and female mice were used for lineage tracing experiment in the Fig. 2. All mice were kept on a 12 h/12 h light/dark cycle under specific pathogen free condition and all animal protocols were approved by the Institutional Animal Experiment Committee of the University of Tsukuba and the Guidelines of Keio University for Animal experiments.

### Cell culture and reagents

Human aortic EC (HAEC; Lonza, CC-2535), human pulmonary artery EC (HPAEC; Lonza, CC-2530) and human cardiac artery EC (HCAEC; Lonza, CC-2585) were grown in EGM™-2 BulletKit™ (for HAEC and HPAEC) and EGM™-2MV BulletKit™ (for HCAEC) supplemented with 2% (v/v) fetal bovine serum and growth factors. These ECs were used between passage 6 and 10. Cobalt chloride (CoCl_2_, 036-03682) was purchased from Wako. VH298, von Hippel-Lindau Tumor Suppressor (VHL) inhibitor (ab230370) was purchased from Abcam. CD45 PTPase inhibitor (#540215) and SB525334 (S1476) were purchased from Merck Millipore and Selleck, respectively. Recombinant human TGFβ1 (100-B-001) and Thrombospondin-1 (3074-TH-050) were purchased from R&D Systems. Matrigel-based hydrogel culture system were employed, and hydrogel concentration and the corresponding Young’s modulus have been previously reported^64^.

### Hypoxia treatment

Stock solution (10 mM) of CoCl_2_ or VH298 was prepared in sterile distilled water and further diluted in medium in order to obtain the final desired concentrations. HAEC, HCAEC and HPAEC were cultured in medium with 100 μM of CoCl_2_ or 100 nM of VH298 with or without 30 nM of CD45 PTPase inhibitor, and incubated for 48 hours (h) in standard CO_2_ incubator.

### Carotid artery ligation

Eight-week-old male C57BL/6 or *Cdh5*-BAC-Cre^ERT2^-LSL-GFP mice were used for carotid artery ligation as previously described^5^. Briefly, the left common carotid artery was dissected free of connective tissues and permanently ligated proximally to the bifurcation. At 1, 2, 3 or 4 weeks after surgery, mice were euthanized, perfused, and right and left carotid arteries were harvested and embedded in optimal cutting temperature compound (OCT, SAKURA Finetek USA Inc.) . Serial cross sections were prepared from the ligature toward the aortic arch with 10 μm intervals until neointima disappeared.

### Immunostaining

Carotid artery was harvested and embedded in OCT and snap frozen in liquid nitrogen. Cross sections of the mouse carotid arteries or HAECs were immediately fixed with 4% paraformaldehyde for 30 min, blocked in 5% normal serum in which secondary antibody was raised, containing 0.1% Tween-20 for 1 h at 37°C. The primary antibodies used are shown in the Extended Data Table 1. Incubation was performed overnight at 4°C. After washing, highly cross-absorbed Alexa 488, Alexa 546 or Alexa 647-conjugated secondary antibodies (Invitrogen) were added at a dilution of 1:200 for 2 h at 37 °C. Control experiments were performed by omitting the primary antibody. Slides were covered with Vectashield containing DAPI (Vector Laboratories) and viewed under a LSM 710 (ZEISS). Colocalization was measured on a single confocal section using the Colocalization-module of Imaris 9.2.1, 64-bit version (Bitplane AG).

### Western blot analysis

HAECs or mouse carotid artery were dissolved in RIPA lysis buffer (Sigma Aldrich, R0278) containing 1% protease inhibitor (Wako, 167-24381) and 1% phosphatase inhibitor (Wako, 160-24371). The lysates were mixed with 3 x SDS sample buffer with 2-mercaptoethanol (Wako, 133-14371) and boiled at 95°C for 5 min, and then were subjected to SDS-PAGE. Proteins were transferred to a PVDF membrane (Millipore, IPVH00010) and immunoblotted with antibodies indicated in the Extended Data Table 1. Membranes were then incubated with respective anti-mouse or anti-rabbit HRP-conjugated secondary antibody (1:1000, Bio-Rad) and visualized with a chemoluminescence kit (Santa Cruz Biotechnology, sc-2048) or SuperSignal West Femto Maximum Sensitivity Substrate (Thermo Fisher Scientific, 34094).

### Fluorescence-activated cell sorting (FACS) analysis

Single cell suspensions from carotid arteries were prepared by enzymatic digestion as previously described^65^. Briefly, right or left carotid aorta were harvested, cut into small pieces and digested in 0.5 ml of enzyme cocktail containing 800 U/mg collagenase type XI (Sigma-Aldrich, C7657), 125 U/mg collagenase type I (Sigma-Aldrich, C0130), 400 U/mg hyaluronidase (Sigma-Aldrich, H3506) and 60 U/mg DNase1 (Sigma-Aldrich, 11284932001) in DPBS with calcium and magnesium (Gibco, 14040-133). Digestion was carried out at 37 °C for 1 h in humid chamber with gently shaking. The cell suspension was filtered through a 100 μm cell strainer (Falcon, 352360) by mashing aorta pieces using a syringe plunger. The cell suspension was centrifuged and resuspended in FACS buffer containing 2% FBS (Gibco) in PBS, then counted. The cell suspension was fixed by 4% PFA for 20 min and washed by FACS buffer, then incubated with fluorescent labelled antibodies, shown in the Extended Data Table 1, for 2 h at 4 °C and washed twice then resuspended in FACS buffer. FACS measurements were performed on FACSAria II™ (BD Biosciences). Data were analyzed using FlowJo software (BD, version 10.7.1).

### Quantitative Polymerase chain reaction (PCR)

RNA was isolated from mouse carotid artery or HAECs using RNeasy Plus Micro Kit (QIAGEN, 74134) and 200 ng of total RNA was subjected to reverse transcription reactions using iScript™ Reverse Transcription Supermix (Bio-Rad, 1708841). SYBR Green was used for amplicon detection and gene expression was normalized to the expression of housekeeping genes Glyceraldehyde 3-phosphate dehydrogenase (GAPDH) or β-actin. PCR reactions were carried out in triplicate in a CFX96 real-time PCR detection system (Bio-Rad) with one cycle of 3 min at 95 °C, then 39 cycles of 10 sec at 95 °C and 30 sec at 55 °C. Levels of mRNA were determined using the ddCt method and expressed relative to the mean dCt of controls. Primer sequences are provided in the Extended Data Table 2.

### Immunoprecipitation

Immunoprecipitation was conducted using a Pierce Crosslink IP kit (Thermo Fisher Scientific, 26147). Antibody against SHARPIN (Proteintech, 14626-1-AP), VE-Cadherin (Santa Cruz Biotechnology, sc-9899) or rabbit normal IgG (10500C, Invitrogen), mouse normal IgG (10400C, Invitrogen) were crosslinked to protein A/G agarose beads and then incubated with protein lysates overnight at 4°C. After washing and elusion, input and IP products were analyzed using Western blotting.

### Atomic Force Microscopy (AFM)

AFM measurements were performed using a NanoWizard IV AFM (Bruker) mounted on top of an inverted optical microscope (IX73, Olympus, Japan) equipped with a digital CMOS camera (Zyla, Andor) as previously described^66^. The AFM quantitative imaging (QI) mode was then used to obtain a force-displacement curve at each pixel of 128 × 128 pixels (100 µm × 100 µm of measured area), using a precisely controlled high-speed indentation test using V-shaped silicon nitride cantilevers with a cone probe (MSNL-10, Bruker) at a spring constant of 0.30 N/m. The QI mode measurements were performed within 1 h after the transfer of the specimen cells to the AFM at room temperature RT (25 °C). These high-speed indentations were performed until reaching a pre-set force of 1 nN. Cell elasticity was calculated from the obtained force-displacement curves by applying a Hertzian model, approximating that the sample is isotropic and linearly elastic. Young’s (elastic) modulus could be extracted by fitting all force-displacement curves with the following Hertzian model approximation:

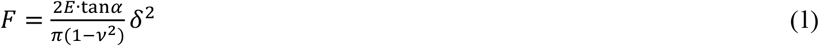

where *F* is the applied force, *E* is the elastic modulus, *ν* is the Poisson’s ratio (0.5 for a non-compressible biological sample), *α* is the opening angle of the cone of the cantilever tip, and *δ* is the indentation depth of the sample. Using the results of the Hertzian model approximation, we identified the Z contact points (specimen surface) and the elastic modulus of the specimens at each pixel, and produced a surface topography map and elastic modulus map of the specimens to estimate cell height and mechanical tension of actin fibers.

### Matrigel tube formation assay

Matrigel (Corning, #356230) was added (300 μl ; 5.45 mg/ml) to each well of a 24-well plate and allowed to polymerize for 1h at 37 °C. 40-50% confluent HAECs were treated with or without 100 μM of CoCl_2_ and/or 30 nM of CD45 PTPase inhibitor for 48 h, then a total 5 × 10^5^ cells were seeded on polymerized-Matrigel. After 8 h, images were obtained by EVOS Imaging system (Thermo Fisher Scientific).

### Statistical analysis

All experiments are presented as means ± standard error of the mean (SEM). Statistical analysis was performed using Prism 8 (Graph Pad, version 8.4.2). A Shapiro-Wilk test was used for the normality test. Statistical significance was determined by either unpaired *t* test, one-way ANOVA, or two-way ANOVA followed by Tukey’s multiple comparison test. If the normality assumption was violated, nonparametric tests were conducted. Mann-Whitney test was used in the Figs. 2e and h, Kruskal-Wallis test with Dunn’s multiple comparisons was used in Fig. 4b (for Thbs1), Fig. 6e (for HIF-2α and pVEC/VEC) and Fig. 7 c (for mean intensity of VE-cadherin), Fig. 7e (for cell height) and Fig. 7f (for Young’s Modulus). *P* < 0.05 denotes statistical significance.

## Author contributions

Y.Y. conceptualized and H.Y. supervised the project. Y.Y. designed and K.R. performed the experiments for carotid artery ligation and analyzed the results. Y.K. provided mice line for lineage tracing and analyzed the data.

Y.Y. and S.T. performed and analyzed in vitro CoCl_2_ and VHL298 experiments. K.N. performed atomic force microscopy analysis. Y.Y. and H.Y. interpreted the data and acquired funding. Y.Y. wrote the manuscript with input from all authors and H.Y. revised the manuscript.

## Acknowledgement

We thank M. Higashi (University of Tsukuba) and A. Hirata (Keio University) for technical assistance. We appreciate M. A. Schwartz (Yale University) and J. Bischoff (Boston Children’s Hospital, Harvard Medical School) for critical reading of the manuscript. We thank Mayumi Mori for assistance with graphics. This work was supported in part by JSPS KAKENHI (Grant Numbers 21H02677), the Japan Foundation for Applied Enzymology and Takeda Science Foundation, the MSD life Science Foundation, SENSHIN Medical Research Foundation, Life Science Foundation, KANAE Foundation for the Promotion of Medical Science, The NOVARTIS Foundation for the Promotion of Science, The MITSUBISHI Foundation and Japan Agency for Medical Research (AMED, Grant Number JP21452398), all to Y.Y. and the JSPS KAKENHI (Grant Number JP17H04289 and JP20H03762) to H.Y.

## Extended Data Figures and Figure legends

**Extended Data Fig. 1.**
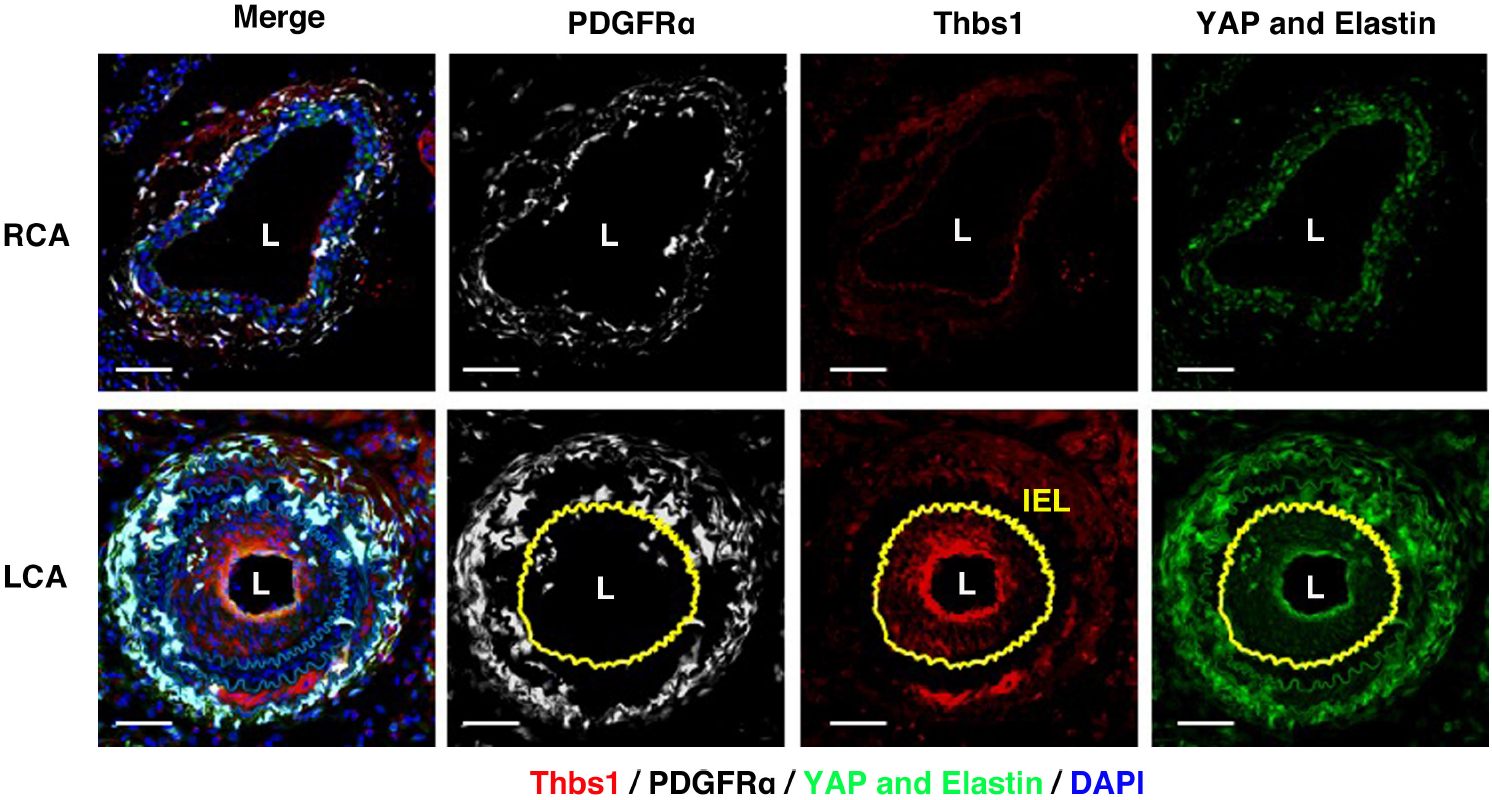
Neointima formation at 4 weeks after carotid artery ligation, related to Fig. 1. Representative immunostaining of cross section of ligated artery (LCA) and contralateral artery (RCA) for PDGFRα (white), Thbs1 (red) and YAP (green) at 4 weeks after ligation in WT mice (n=3). DAPI (blue) and auto-fluorescence of elastin (green). Yellow line shows internal elastin lamina (IEL). L: lumen. Bars are equal 50 μm.

**Extended Data Fig. 2.**
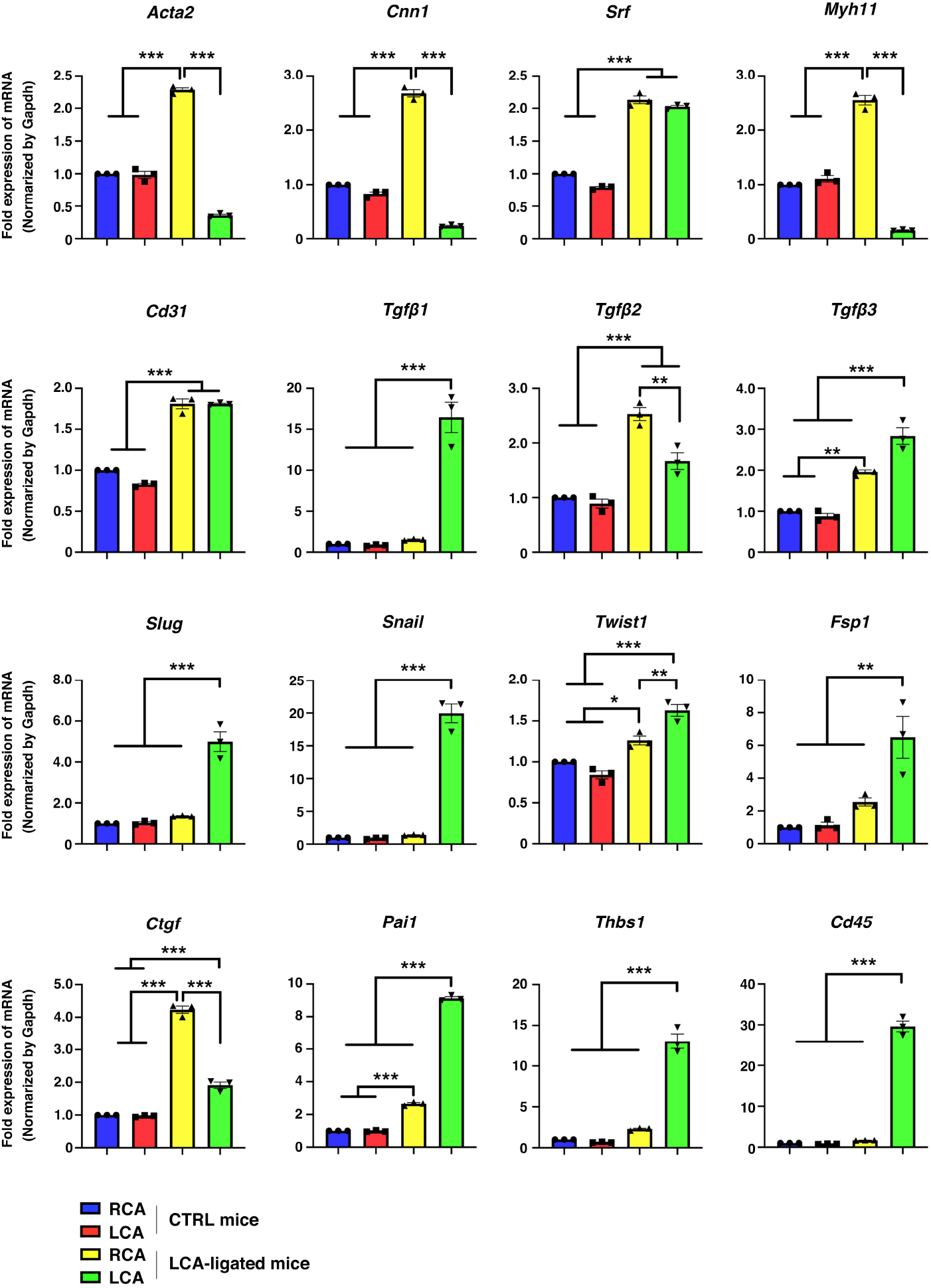
Gene expression profiles on carotid artery, related to Fig. 4a. qPCR analysis of SMC contractile genes, EndMT markers, and TGFβ signal genes from right carotid artery (RCA) or left carotid artery (LCA) of CTRL (n=3) and LCA-ligated mice (n=3). Bars indicate mean ± SEM. \*\*\**P*< 0.001, \*\**P*< 0.01, one-way ANOVA.

**Extended Data Fig. 3.**
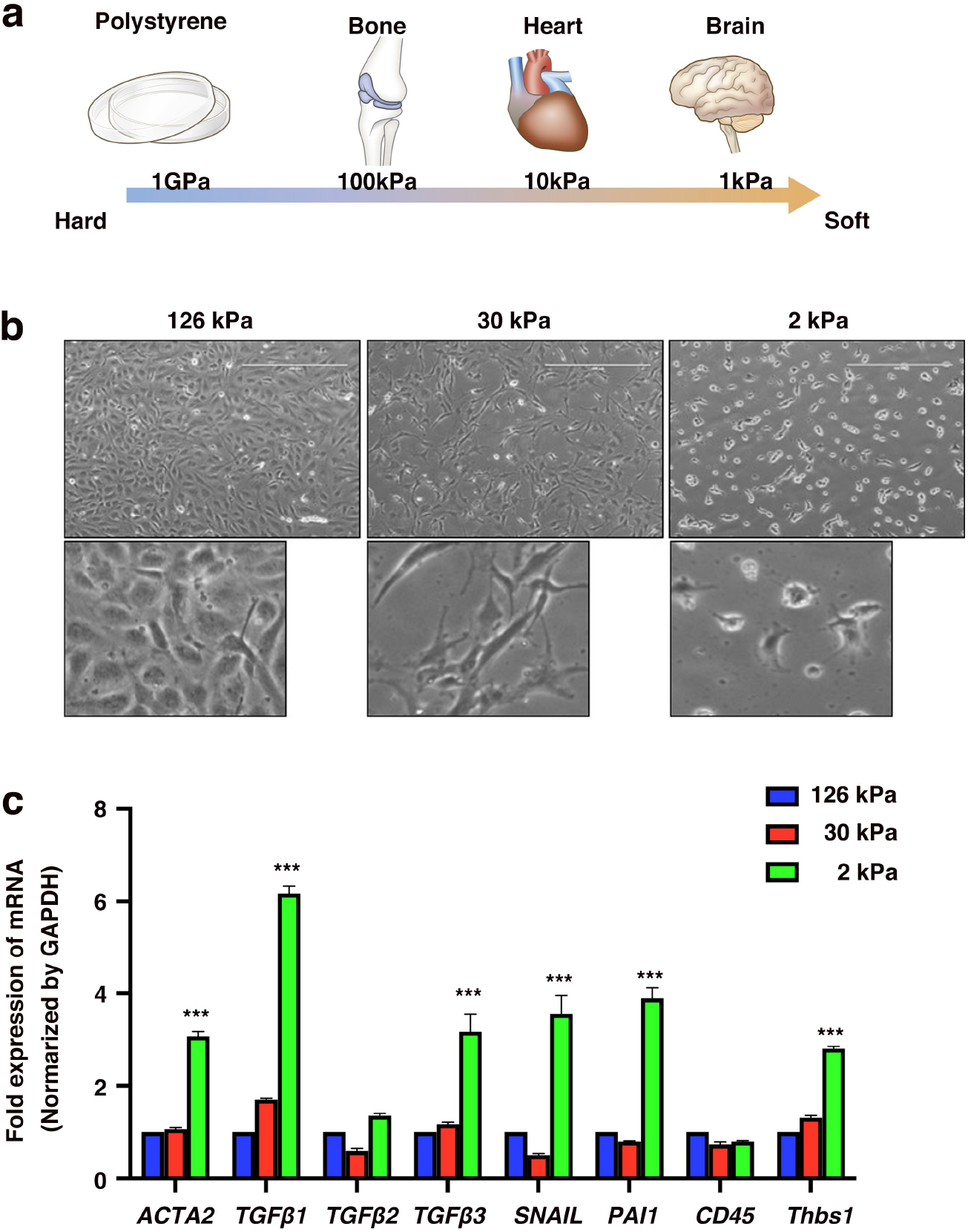
Soft matrix promoted EndMT but not induce CD45 expression, related to Fig. 5a. **(a)** Illustrating the range of stiffness in the solid tissues and polystyrene dish or hydrogels (126 kPa, 30 kPa and 2 kPa). **(b)** Bright field-images of HAECs seeded after 24 h on different stiffness of hydrogels. (**c)** qPCR analysis of SMC contractile gene, *ACTA2* and EndMT markers, and TGFβ signal genes from HAECs on different stiffness of hydrogels. n=3 for each condition. Bars indicate mean ± SEM. \*\*\**P*< 0.001, one-way ANOVA.

**Extended Data Fig. 4.**
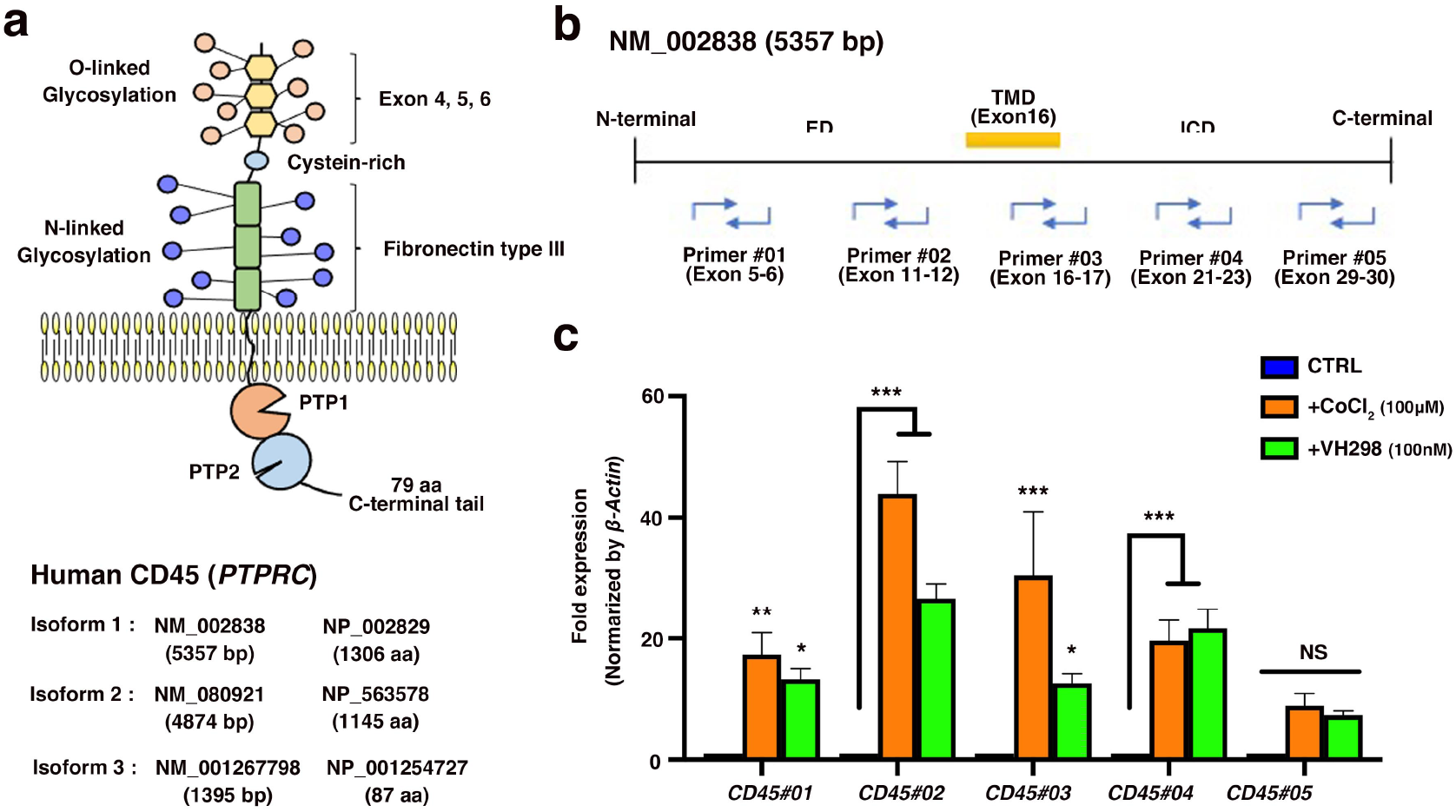
Evaluation of CD45 expression by qPCR, related to Fig. 5a. **(a)** Illustrating a structure of CD45 and genomic information for human CD45. **(b)** Primer design for qPCR analysis. ED: extracellular domain, TMD: trans-membrane domain, ICD: intracellular domain. **(c)** qPCR analysis of CD45 using different primer-sets. HAECs were cultured with CoCl_2_ (100 μM) or VH298 (100 nM) for 48 h. n=3 for each condition. Bars indicate mean ± SEM. \**P*< 0.05, \*\**P*< 0.01, \*\*\**P*< 0.001, two-way ANOVA. Following experiments, we used Primer_01 (*CD45#01*) to test mRNA expression of CD45.

**Extended Data Fig. 5.**
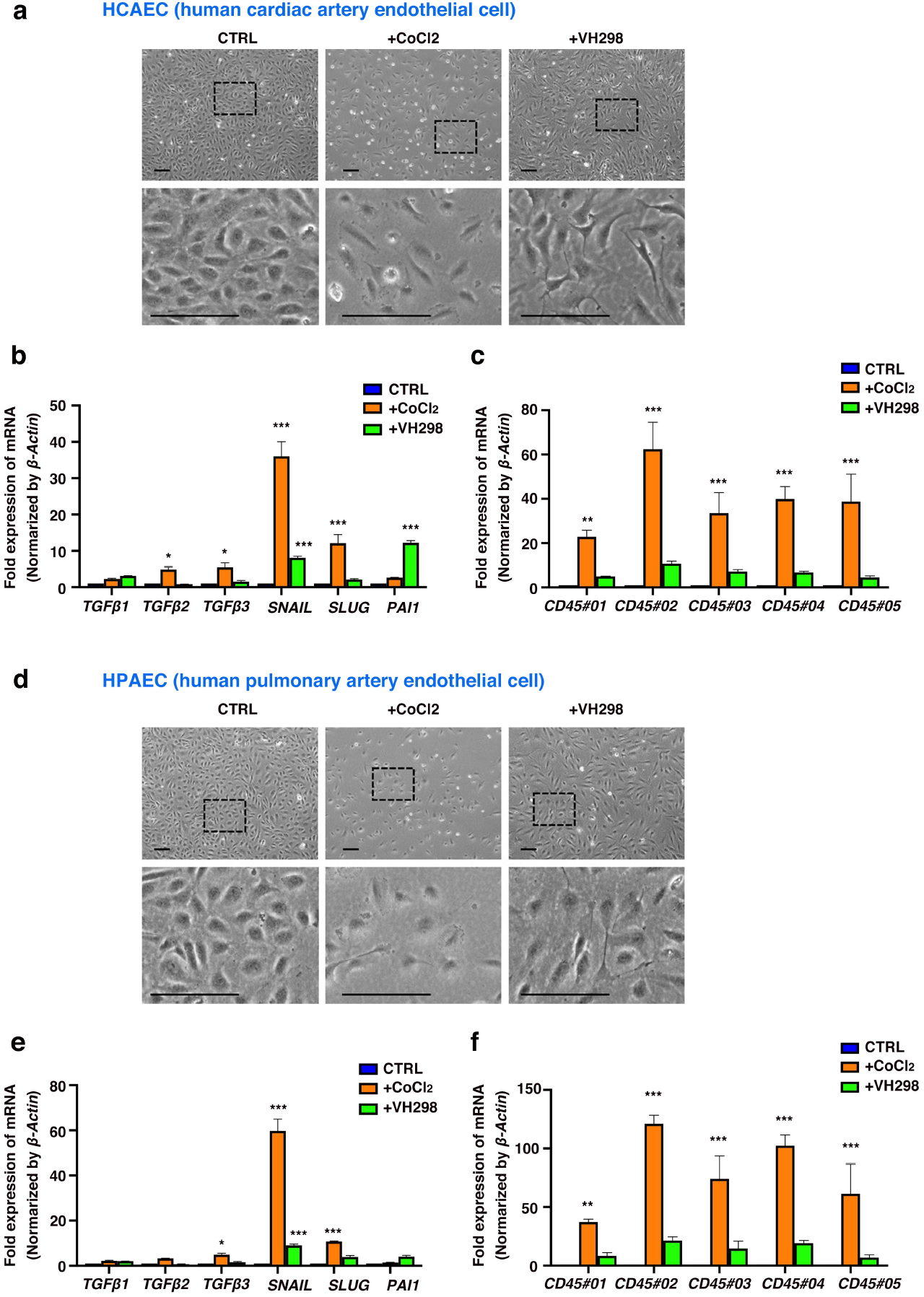
Evaluation of CD45 and EndMT marker expression on HCAEC and HPAEC, related to Fig. 5a. **(a and d)** Cell morphology after 48 h CoCl_2_ (100 μM) or VH298 (100 nM) treatment. **(b, c and e, f)** qPCR analysis of EndMT markers (b and e) and CD45 using different primer-sets (c and f). a-c for HCAEC, d-f for HPAEC. n=3 per conditions. Bars indicate mean ± SEM. \**P*< 0.05, \*\**P*< 0.01, \*\*\**P*< 0.001, two-way ANOVA.

**Extended Data Fig. 6.**
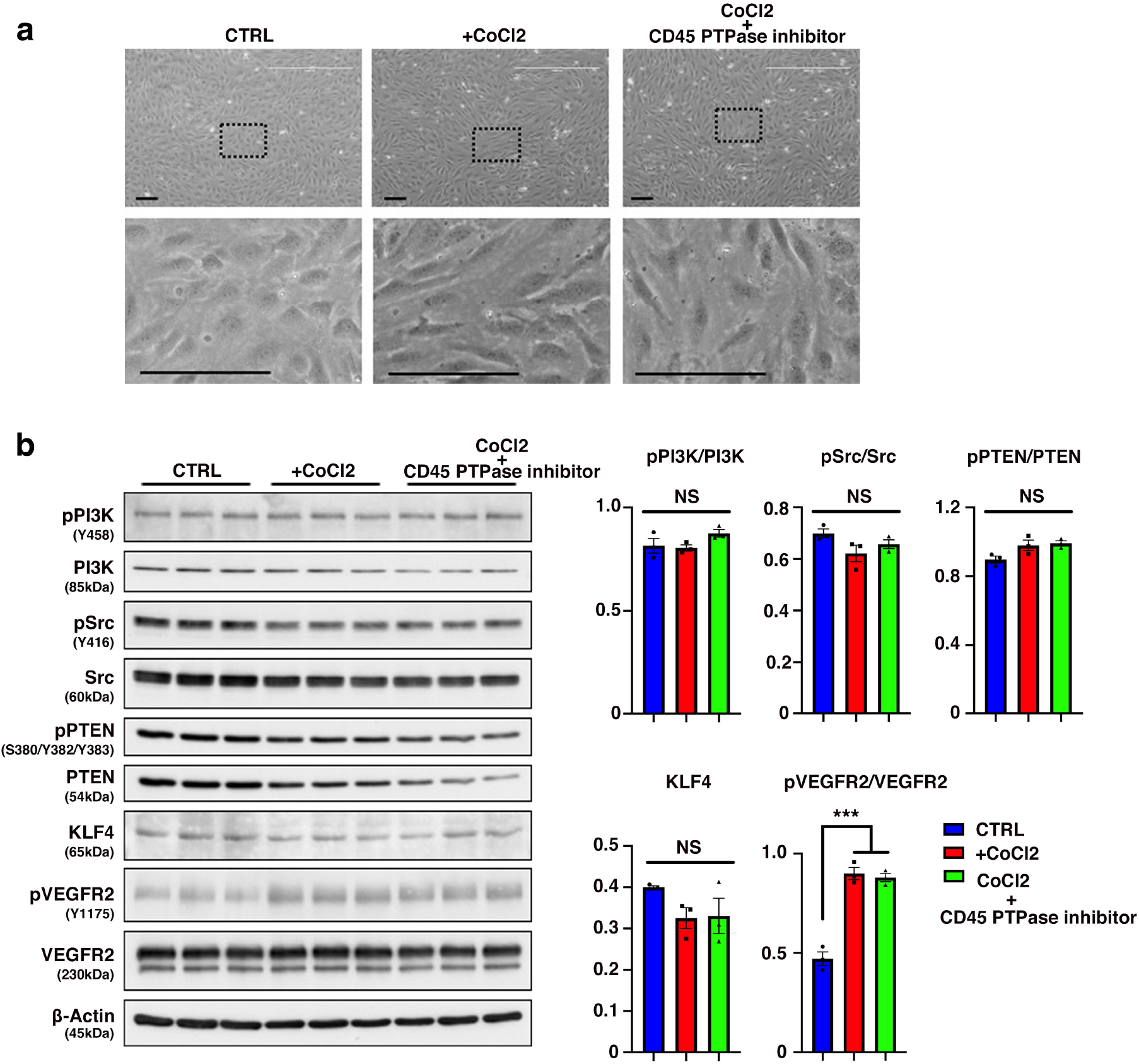
Effects on CD45 PTPase inhibitor under CoCl_*2*_ treatment, related to Fig. 6e. **(a)** Cell morphology under CoCl_2_ (100 μM) with or without CD45 PTPase inhibitor (30 nM) treatment. **(b)** Western blots show PI3K, Src, PTEN KLF4 and VEGFR2 levels in CTRL, CoCl_2_ (100 μM) and CoCl_2_ with CD45 PTPase inhibitor (30 nM). n=3 for each condition. Quantification graphs are shown on the right. Bars indicate mean ± SEM. \*\*\**P*< 0.001, one-way ANOVA. β-actin blot is same as shown in Fig. 6e.

**Extended Data Table 1.** Antibodies used in this study Western blot analysis.

| **Antibody** | **Dilution** | **Source** | **Catalog number** |
| --- | --- | --- | --- |
| **Thbs1** | 1:1000 | NeoMarkers | MS-421 |
| **pYAP (S127)** | 1:500 | Cell Signaling | #4911 |
| **YAP** | 1:1000 | Cell Signaling | #14074 |
| **pSmad2 (S465/S467)** | 1:500 | Cell Signaling | #3108 |
| **Smad2/3** | 1:1000 | Cell Signaling | #8685 |
| **Slug** | 1:500 | Cell Signaling | #9585 |
| **pERK1/2 (T202/Y204)** | 1:500 | Cell Signaling | #4376 |
| **ERK1/2** | 1:500 | Cell Signaling | #9102 |
| **GAPDH** | 1:1000 | Cell Signaling | #2118 |
| **HIF-1α** | 1:1000 | Cell Signaling | #36169 |
| **HIF-2α** | 1:1000 | Cell Signaling | #7096 |
| **pVE-Cadherin (Y658)** | 1:500 | Invitrogen | 44-1144G |
| **VE-Cadherin** | 1:1000 | Invitrogen | 36-1900 |
| **pVEGFR2 (Y1175)** | 1:1000 | Cell Signaling | #3770 |
| **VEGFR2** | 1:1000 | Cell Signaling | #9698 |
| **Integrin α11** | 1:1000 | Abcam | Ab198826 |
| **Integrin αv** | 1:500 | Cell Signaling | #4711 |
| **Integrin β1** | 1:500 | Cell Signaling | #4706 |
| **pAKT (Y308)** | 1:500 | Cell Signaling | #13038 |
| **AKT** | 1:1000 | Cell Signaling | #9272 |
| **SHARPIN** | 1:500 | Proteintech | 14626-1-AP |
| **pPI3K (Y458)** | 1:500 | Cell Signaling | #4228 |
| **PI3K** | 1:500 | Cell Signaling | #4257 |
| **pSrc (Y416)** | 1:500 | Cell Signaling | #6943 |
| **Src** | 1:500 | Cell Signaling | #2109 |
| **pPTEN (S380/T382/T383)** | 1:500 | Cell Signaling | #9549 |
| **PTEN** | 1:500 | Cell Signaling | #9188 |
| **KLF4** | 1:500 | Cell Signaling | #4038 |
| **Fibronectin** | 1:500 | BD Biosciences | 610078 |
| **Collagen I** | 1:500 | Merck (Millipore) | AB765P |
| **KDEL (BiP)** | 1:500 | Enzo Life Sciences | ADI-SPA-827 |
| **p120 Catenin** | 1:500 | Cell Signaling | #59854 |
| **β-Catenin** | 1:500 | Cell Signaling | #19807 |
| **β-Actin** | 1:1000 | Cell Signaling | #3700 |

Immunostaining
| **Antibody** | **Dilution** | **Source** | **Catalog number** |
| --- | --- | --- | --- |
| **αSMA (1A4)** | 1:200 | Sigma-Aldrich | A2547 |
| **CD31** | 1:200 | BD Pharmingen | 550274 |
| **SM-MYH** | 1:200 | Alfa Aesar | J64817 |
| **PECAM (M-20)** | 1:200 | Santa Cruz Biotechnology | sc-1506 |
| **CD68** | 1:200 | Santa Cruz Biotechnology | sc-20060 |
| **CD11b (integrin αM)** | 1:200 | Santa Cruz Biotechnology | sc-53086 |
| **VE-Cadherin (F-8)** | 1:200 | Santa Cruz Biotechnology | sc-9989 |
| **CD45 (PTPRC)** | 1:200 | BD Pharmingen | 550539 |
| **FSP-1 (S100A4)** | 1:200 | NeoMarkers | RB-1804-A0 |
| **pSmad2/3 (S423/S425)** | 1:200 | Santa Cruz Biotechnology | sc-11769-R |
| **HIF-1α** | 1:200 | Santa Cruz Biotechnology | sc-13515 |
| **VE-cadherin** | 1:500 | Invitrogen | 36-1900 |
| **Rhodamine Phalloidin** | 1:500 | Cytoskeleton | #PHDR1 |
| **YAP** | 1:200 | Cell Signaling | #14074 |
| **Thbs1** | 1:200 | NeoMarkers | MS-421 |
| **PDGFRα** | 1:200 | R&D Systems | AF1062 |

FACS
| **Antibody** | **Dilution** | **Source** | **Catalog number** |
| --- | --- | --- | --- |
| **αSMA-PE** | 1:200 | R&D Systems | IC 1420P |
| **CD31-APC** | 1:200 | Invitrogen | 17-0311-82 |
| **CD45-FITC** | 1:200 | Invitrogen | 11-0451-85 |

Immunoprecipitation
| **Antibody** | **Amount** | **Source** | **Catalog number** |
| --- | --- | --- | --- |
| **SHARPIN** | 3.27μg | Proteintech | 14626-1-AP |
| **VE-Cadherin** | 4.00μg | Santa Cruz Biotechnology | Sc-9989 |
| **Rabbit control IgG** | 1.5µg | Invitrogen | C10500 |
| **Mouse control IgG** | 3.0µg | Invitrogen | C10400 |

**Extended Data Table 2.**
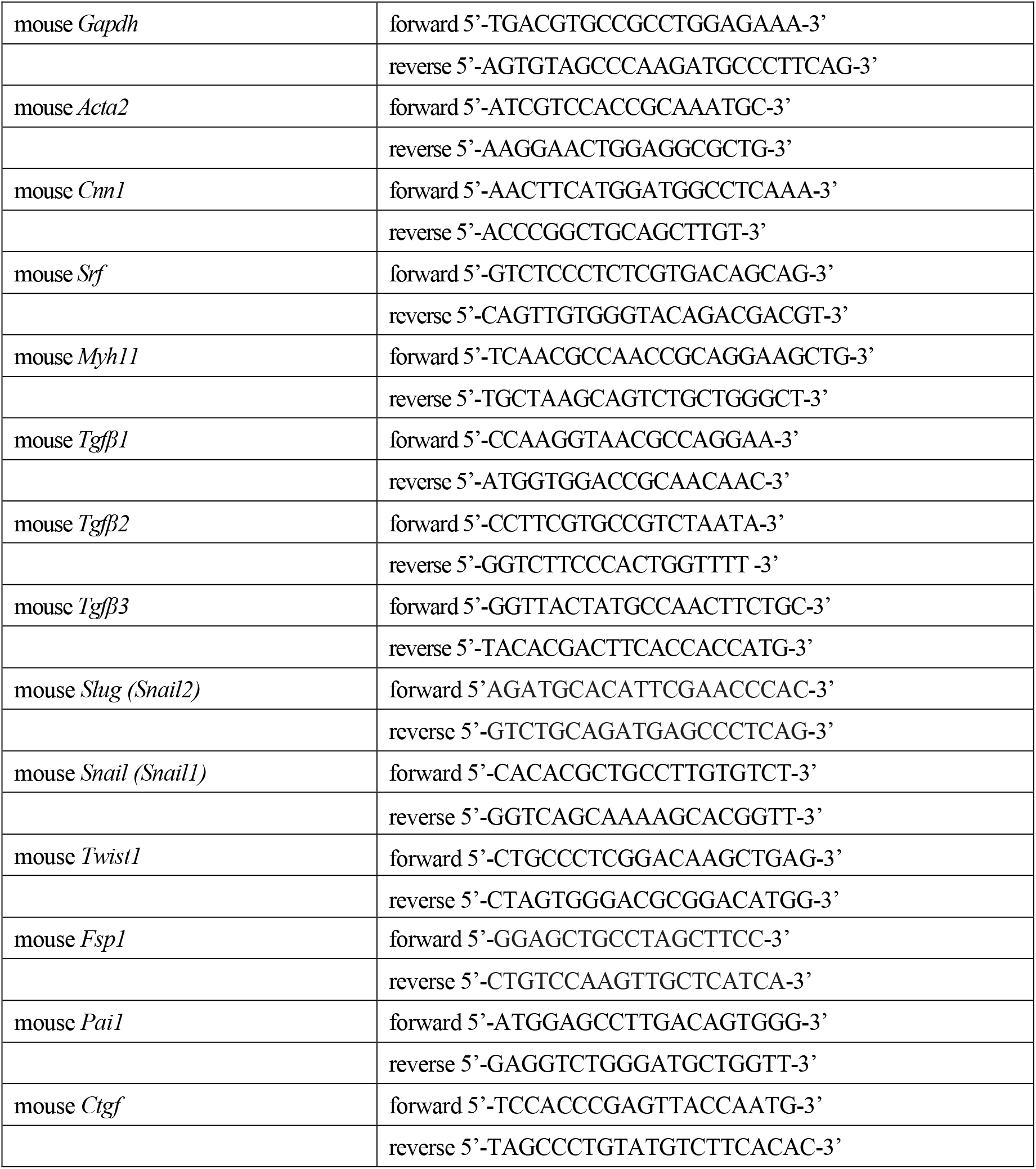
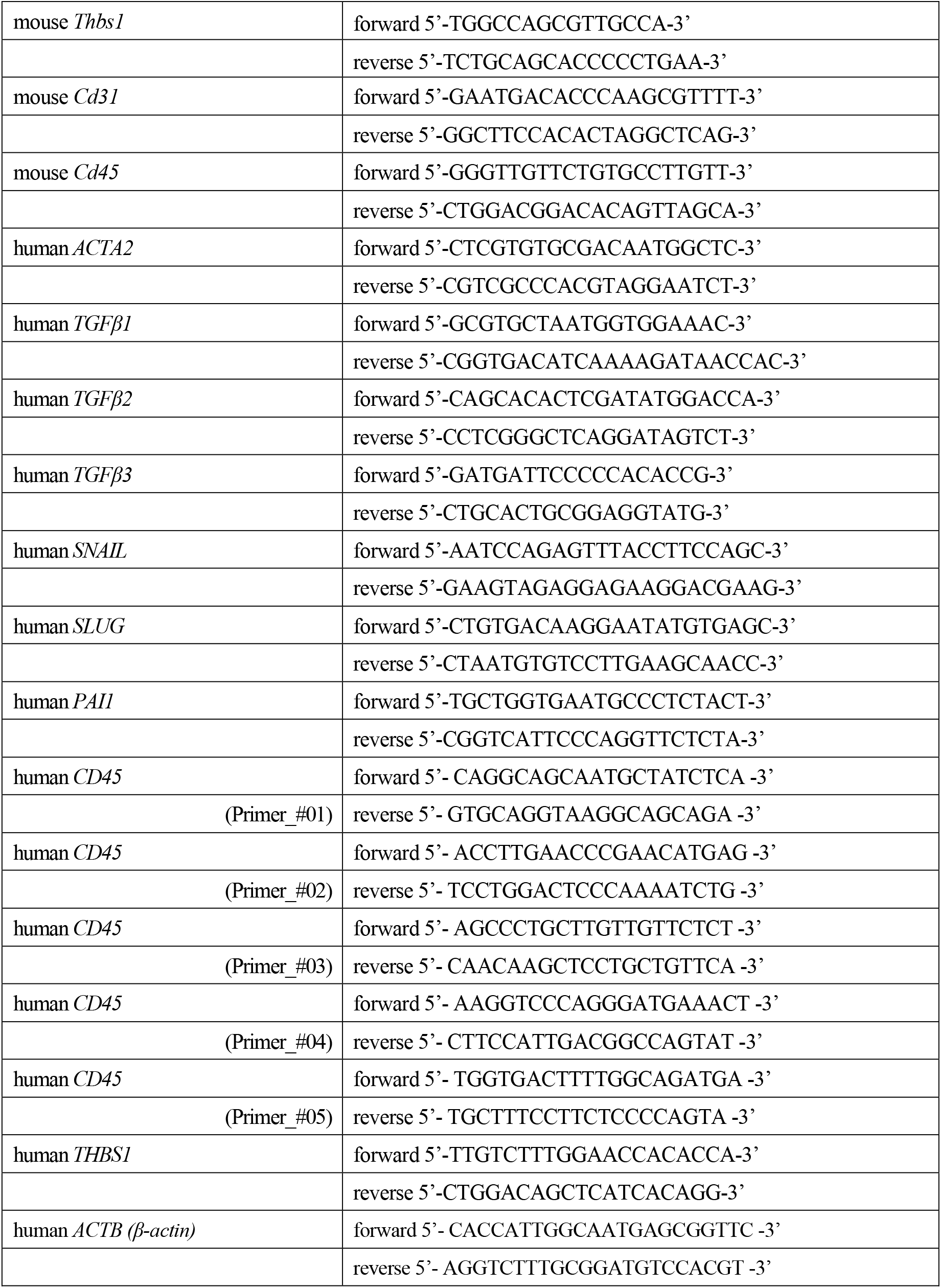
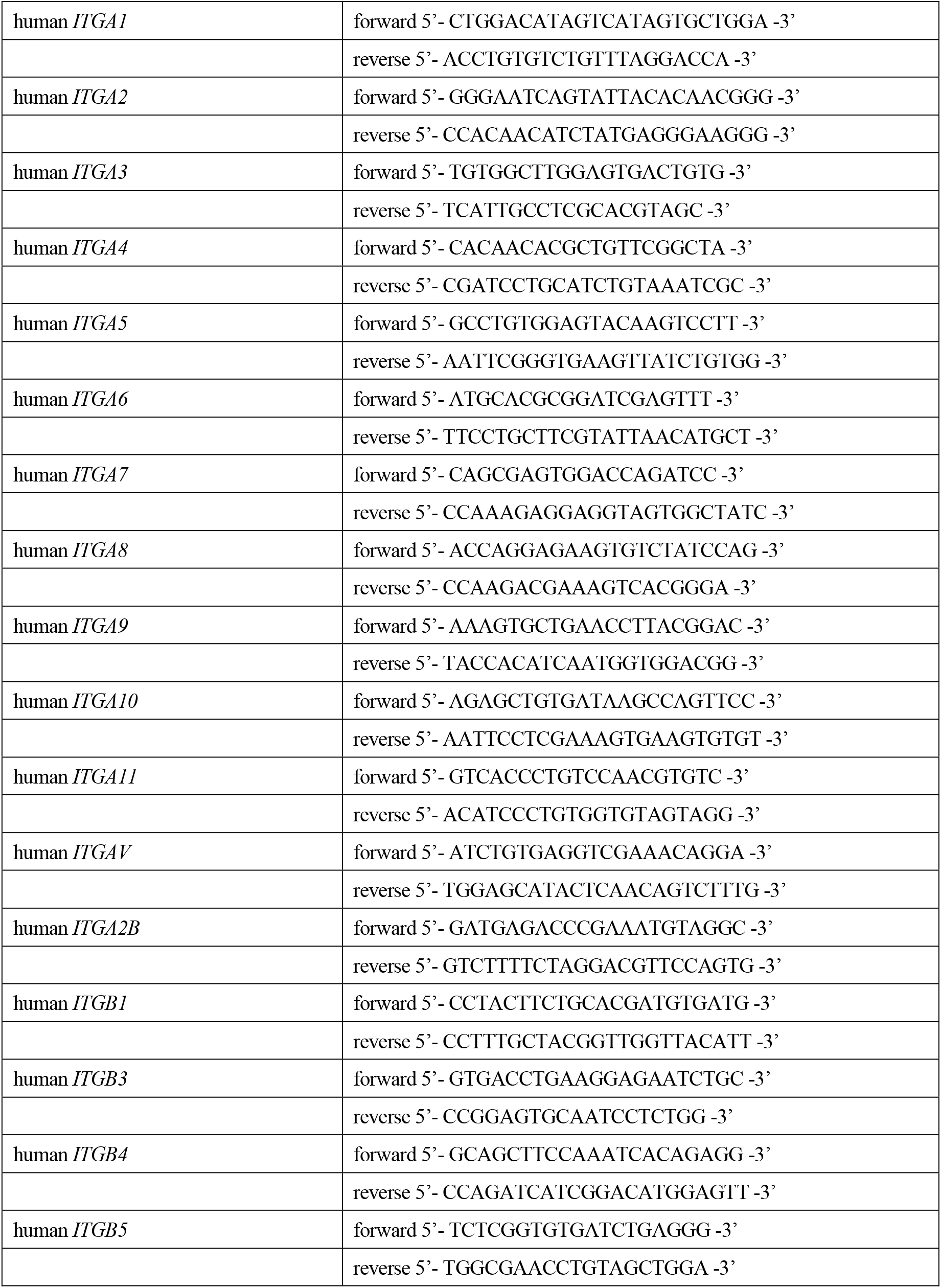
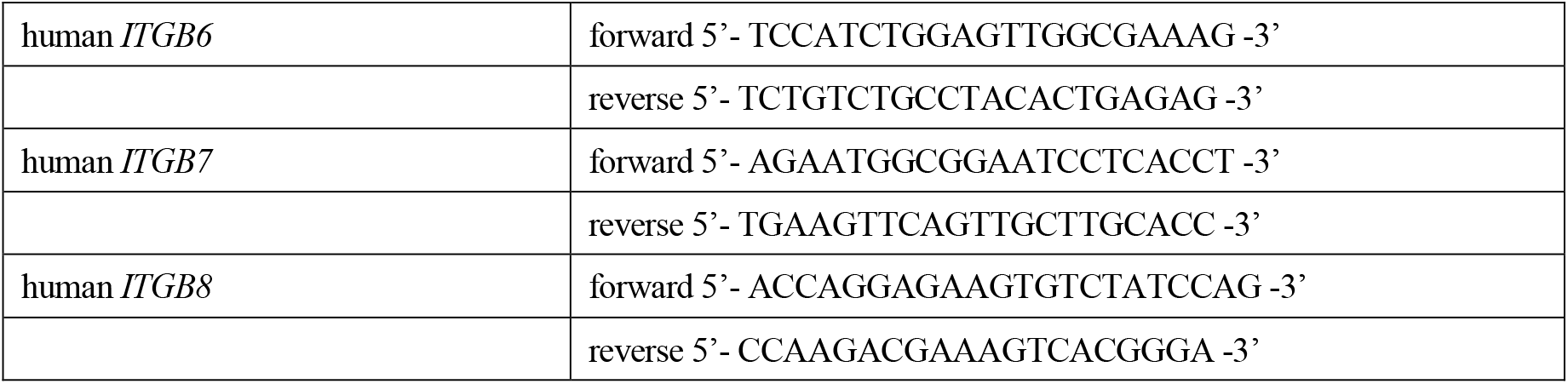
qPCR primer sequences.

